# New cerebellar nuclei evolve through diversification of excitatory progenitors

**DOI:** 10.64898/2026.09.17.751883

**Authors:** Manjari M-G Anant, Eli Clemens Zuercher, Maggie Lowman, Caleb Shi, Dylan Z. Faltine-Gonzalez, Michael L. Piacentino, Jean Fan, Justus M. Kebschull

## Abstract

New brain regions have been proposed to evolve via the duplication of an existing region’s complete set of cell types. Here we investigate the mechanism underlying such apparent evolutionary duplications in the cerebellar nuclei, the output regions of the cerebellum. Using developmental single-cell and spatial transcriptomics time courses in mouse and chicken, we show that formation of new nuclei is governed by changes to nucleus-specific excitatory progenitors. During development, excitatory neurons transcriptomically and spatially define cerebellar nuclei territories, with nucleus- and species-invariant inhibitory neurons incorporating into these territories only after they are established. Across species, evolutionarily conserved excitatory neuron progenitors can generate new nuclei by co-opting border formation gene programs to spatially sort their descendant neurons into distinct territories. While most new nuclei share a common developmental origin and the canonical olivocerebellar error-prediction circuitry, we find the higher order subnuclei of the ancient medial nucleus have a distinct origin and do not participate in the canonical circuit, suggesting a unique evolutionary innovation. Together, our findings suggest that new brain regions can evolve when conserved excitatory cell types first diversify and then spatially segregate, while generic inhibitory neurons are incorporated after regions have formed.

## MAIN

A major unresolved question in the evolution of the nervous system is how new brain regions with all their constituent cell types arise over evolutionary time. One influential model proposes that an ancestral cell type set corresponding to a complete region can duplicate as a whole and then diverge to take on new functions, similar to gene duplication and divergence during genome evolution^1–8^. However, while this duplication and divergence model explains the adult state of the brain, it lacks any mechanistic insight into how such a duplication may be implemented during development. Region duplication could be supported by diverse developmental mechanisms^8^. For example, new regions might form via the generation of new region-specific progenitors, through spatial segregation of conserved ancestral cell types, or by a newly evolved external region-patterning event to impose region boundaries on similar neurons. Each potential mechanism has different implications on how easy it is to evolve a new brain region, how the formation of new regions contributes to new brain functions, and how resilient regions are to changes in brain size or to disease.

The cerebellar nuclei, the sole output structures of the cerebellum, provide an ideal system to dissect the mechanisms of region duplication. The number of cerebellar nuclei has increased over evolutionary time from a putative single ancestral nucleus still found in cartilaginous fishes and amphibians, to two in birds and reptiles, and three or four in mammals^7^ (**Fig. 1A**). Despite these differences in numbers, each cerebellar nucleus of birds and mammals is composed of a copy of a conserved, archetypal cell type set, thus suggesting that the cerebellar nuclei evolved by a repeated process of duplication and divergence^3^.

**Figure 1.**
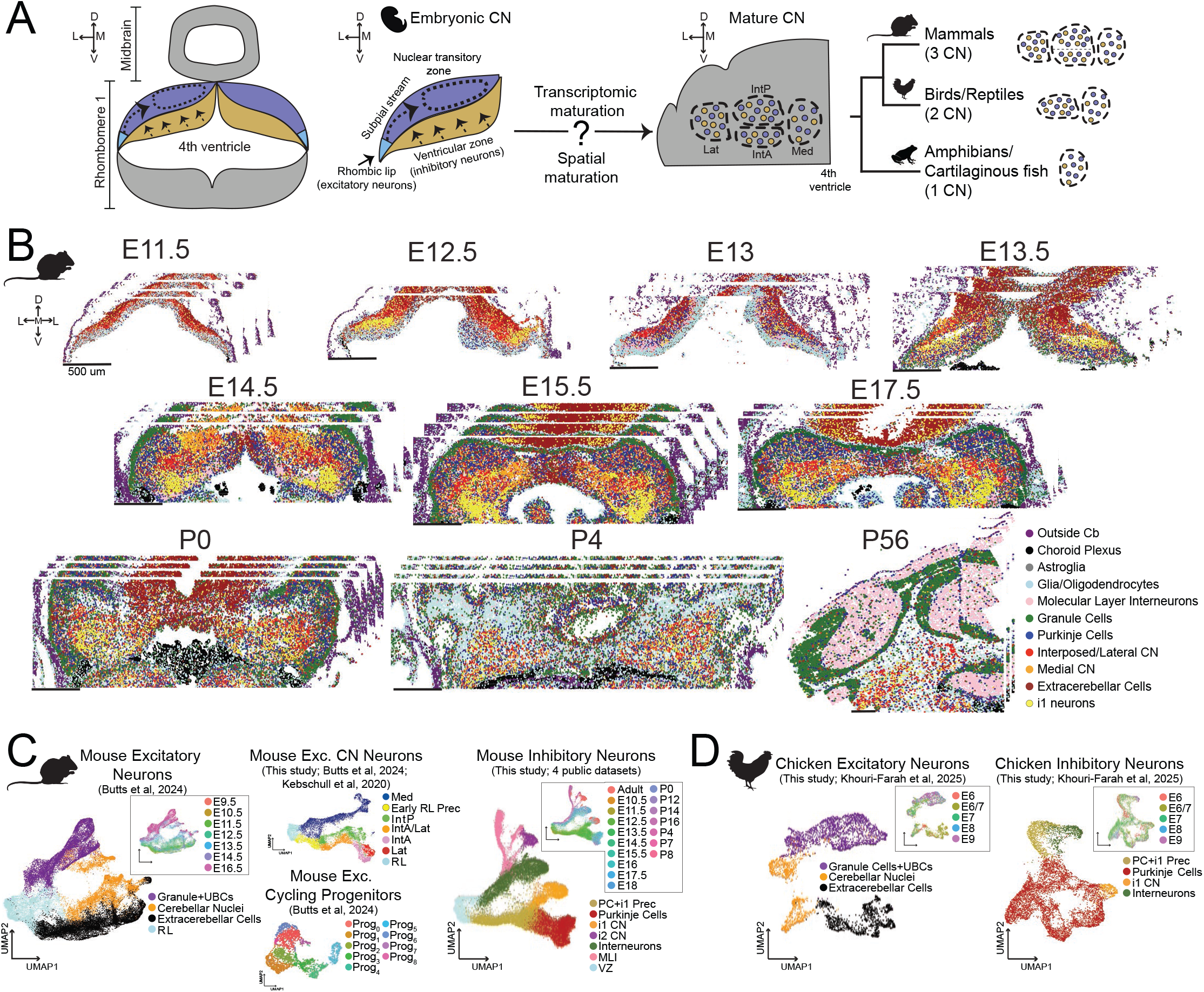
Single-cell and spatial transcriptomic atlases of cerebellar nuclei development in mouse and chicken. **(A)** Schematic of the known developmental migration patterns and adult arrangement of cerebellar nuclei across species, and the developmental and evolutionary questions addressed in this study. **(B)** BARseq3 spatial transcriptomic data from embryonic day (E) 11.5 to postnatal day (P) 56, colored by integrated cell type identity. N=33 sections. **(C)** UMAP embeddings of the mouse scRNAseq datasets used in this study, colored by cell type and, in insets, timepoint. **(D)** UMAP embeddings of the chicken scRNAseq datasets used in this study, colored by cell type and, in insets, timepoint.

The cerebellar nuclei also provide an opportunity to examine how new brain regions acquire distinct functions. In mammals, the cerebellar nuclei are divided into the medial (fastigial), anterior and posterior interposed (IntP, IntA), and lateral (dentate) nuclei. Each nucleus is functionally specialized, with phylogenetically older, medial nuclei generally performing brainstem and vestibular functions and phylogenetically younger, lateral nuclei taking on higher-order functions^7,9,10^. However, recent evidence of higher-order function in the putatively oldest medial nucleus complicates this simple picture^11–16^. Resolving how new nuclei are formed during development and evolution thus may help explain how the diverse functions of the cerebellar nuclei emerged.

Currently, little is known about how individual cerebellar nuclei develop, when and how new nuclear identities are established, and how developmental programs vary across species to generate different numbers of cerebellar nuclei (**Fig. 1A**). Here we use large scale BARseq3 spatial transcriptomics^17^ and single-cell/nucleus RNA sequencing (scRNAseq) in a developmental time course to understand cerebellar nuclei development and interrogate the mechanisms underlying cerebellar nuclei duplication in mouse and chicken. We find that region formation is hierarchically organized, with excitatory neurons born from region-specific progenitors that establish nuclear territories, whereas nucleus-invariant inhibitory neurons fill in later during development. Within this scheme, all canonical cerebellar nuclei are born from the early rhombic lip in a lateral−to−medial, higher-order−to−basal-function order. Progenitors for new mammalian specific nuclei already exist in chickens, but uniquely co-opt RA-FGF8 boundary formation programs to drive the emergence of a new nucleus in mammals. The only exception to this rule is the higher-order subnuclei of the medial nucleus, which derive from the late rhombic lip and form an alternative cerebellar circuit, thus likely having a distinct evolutionary origin.

## RESULTS

### A spatial transcriptomics atlas of cerebellar nuclei development

The cerebellar nuclei are the first-born neurons of the cerebellum and arise from two germinal zones: excitatory cerebellar nuclei neurons are born at the upper rhombic lip (RL)^18–20^, while inhibitory neurons are born from the ventricular zone lining the fourth ventricle^21–23^ (**Fig. 1A**). Postmitotic excitatory neurons migrate tangentially along the subpial stream to reach the nuclear transitory zone, where they mix with radially migrating inhibitory neurons to eventually form the adult cerebellar nuclei^24–26^. To understand how these two cell populations form new cerebellar nuclei in space and time, we collected a comprehensive spatial transcriptomics time course of mouse cerebellar nuclei development. For optimal coverage and ability for discovery, we used BARseq3^17^, a recently developed high-resolution spatial transcriptomics method, to measure the expression of an unbiased gene panel of 1,761 genes, that contained essentially all mouse transcription factors and cell adhesion/migration molecules, as well as neuronal marker genes (**Extended Data Table 1**). We generated datasets from coronal cerebellar sections at 10 timepoints spanning E11.5 to adulthood (E11.5, E12.5, E13, E13.5, E14.5, E15.5, E17.5, P0, P4, and P56), sampling 2 to 4 anterior-posterior positions at each timepoint. In total, we measured 33 sections with 998,978 cerebellar cells (**Fig. 1B**).

We further complemented this high spatial resolution but gene set restricted view of cerebellar nuclei development with a full-transcriptome scRNAseq time-course of cerebellar nuclei development in mice by combining self-generated and publicly available scRNAseq datasets spanning E9.5 through adult stages (**Fig. 1C; Supplementary Fig. 2A-F**)^3,27–30^. To enable evolutionary comparisons, we also assembled a comparable chicken dataset from self-generated and public data^27^ spanning from E6 to E9 (corresponding to ∼E13.5 to E15.5 in mice^31^; **Fig. 1D; Supplementary Fig. 2G-J**).

### UMAP1

In both BARseq3 and scRNAseq datasets, we recovered the expected cell types and spatial gene expression patterns. Excitatory lineages contained extracerebellar-fated neurons, cerebellar nucleus neurons, unipolar brush cells and granule cells, appearing in the expected developmental sequence and expressing established markers (**Extended Data Figs. 1; 2A,B,E,F; Supplementary Fig. 1B**)^30,32–34^. Inhibitory lineage embeddings similarly resolved cerebellar cortical and nuclear interneurons, Purkinje neurons, and i1 cerebellar nuclei neurons with expected marker genes and developmental timing (**Extended Data Figs. 1; 2D-F; Supplementary Fig. 1B**)^27,35^. Together these spatial and single-cell data provide a wholistic view of cerebellar nucleus cell identity, developmental state, and spatial organization that form the basis for this study.

All curated datasets are available to browse in an interactive format at https://mmganant.github.io/CNdev_website/.

### New excitatory cerebellar nuclei neurons derive from the early rhombic lip

To understand how cerebellar nuclei formation is implemented during development and evolution, we asked whether the excitatory or inhibitory neurons of the cerebellar nuclei, or both, drive changes in nuclear identity and number. We first focused on the excitatory neurons (**Figs. 2-4**) and return to inhibitory neurons in **Fig. 5**.

**Figure 2.**
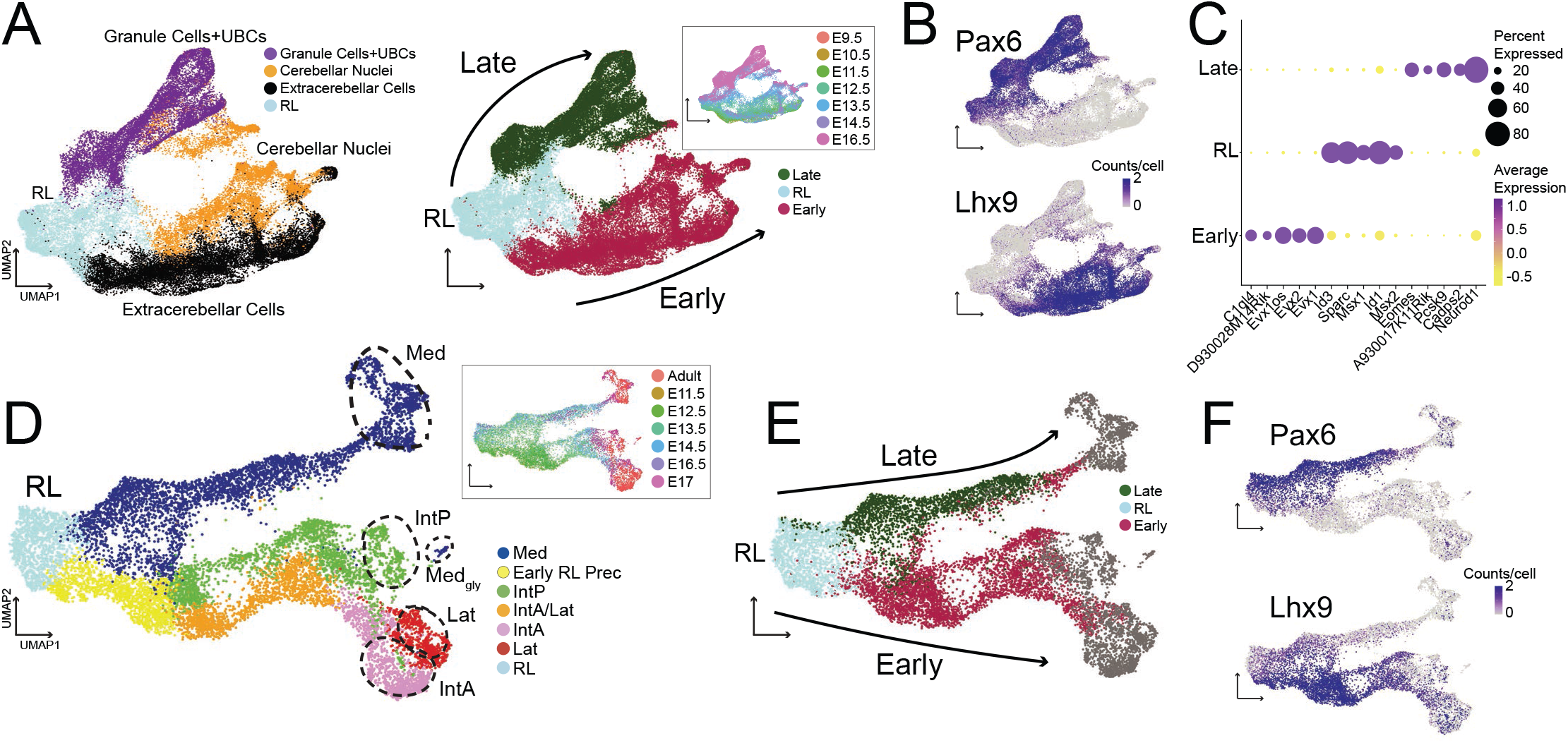
Cerebellar nuclei derive from both early and late rhombic lip. **(A)** UMAP embedding of RL-derived mouse excitatory neurons reveals two major branches corresponding to early and late RL waves. **(B)** Expression of marker genes distinguishing early (*Lhx9+*) and late branches (*Pax6+*). **(C)** Dot plot of differentially expressed genes between early RL, late RL, and early postmitotic RL neurons (RL). **(D)** UMAP embedding of excitatory cerebellar nucleus neurons comprising developmental data from ^40^, additional E17.5 data, and the ^3^ adult dataset. Adult nucleus labels from ^3^ are indicated. Med, whole medial nucleus; Medgly, large, RL-derived glycinergic neurons in medial nucleus; Lat, lateral nucleus. **(E)** Projection of early and late rhombic lip labels from **(A)** onto the integrated excitatory cerebellar nucleus dataset. **(F)** Expression of marker genes distinguishing early (*Lhx9+*) and late branches (*Pax6+*) in the excitatory cerebellar nucleus neuron dataset.

**Figure 3.**
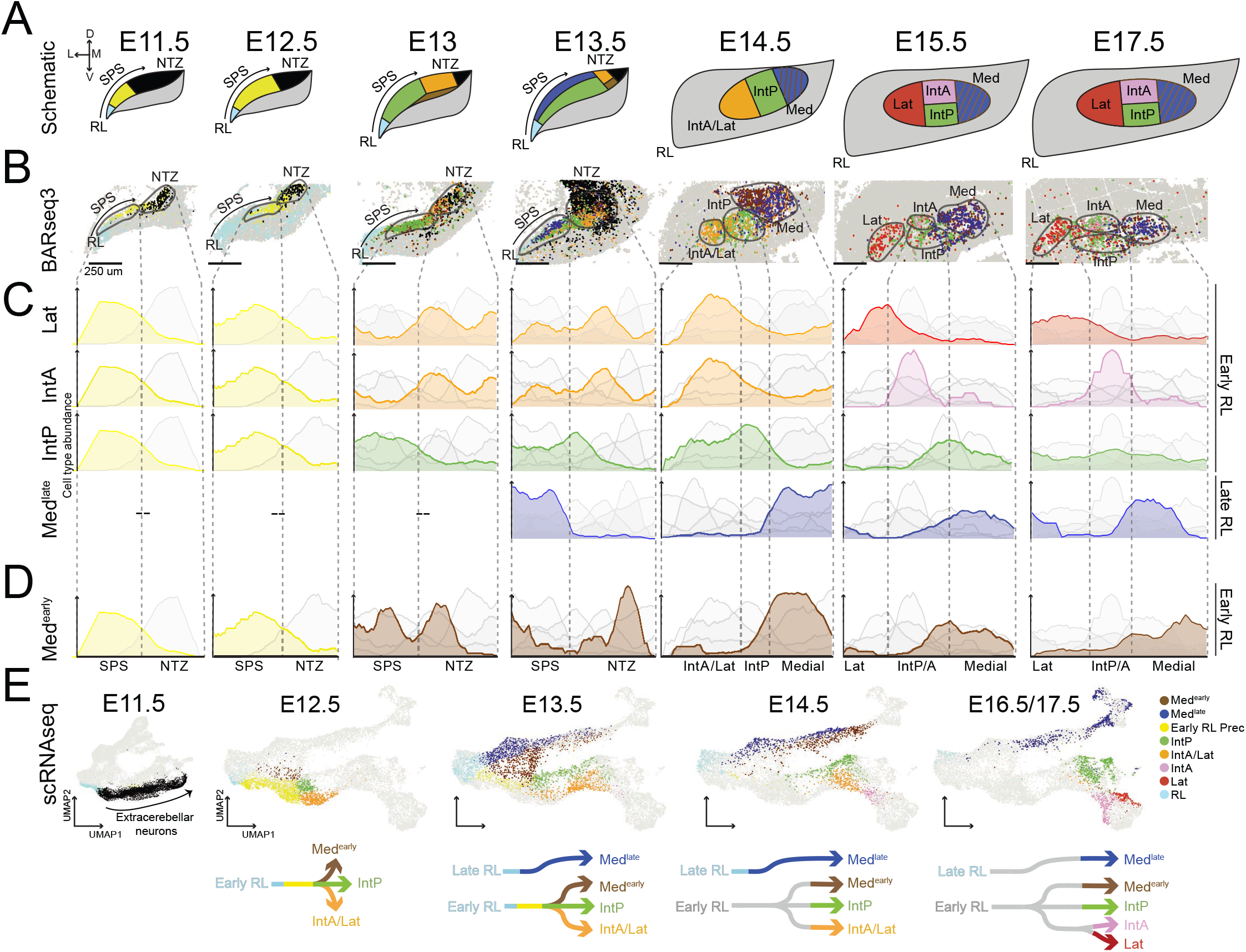
Excitatory neurons are nucleus-specific before forming nuclear territories. **(A)** Schematic summarizing the migration patterns of excitatory cerebellar nucleus cell types during development in mouse. SPS, subpial stream; NTZ, nuclear transitory zone; Med, medial nucleus, Lat, lateral nucleus. **(B)** Representative BARseq3 slices showing one hemisphere of the cerebellar anlage at indicated timepoints, with excitatory neurons colored by cell type. **(C,D)** Quantification of cell type abundances along the demarcated mediolateral axis for populations shown in **(B)**. **(E)** Integrated scRNAseq embedding of cerebellar nucleus excitatory neurons demonstrating close correspondence with BARseq3 spatial data at matched developmental timepoints. Arrows summarize UMAP trajectories at the indicated timepoint.

**Figure 4.**
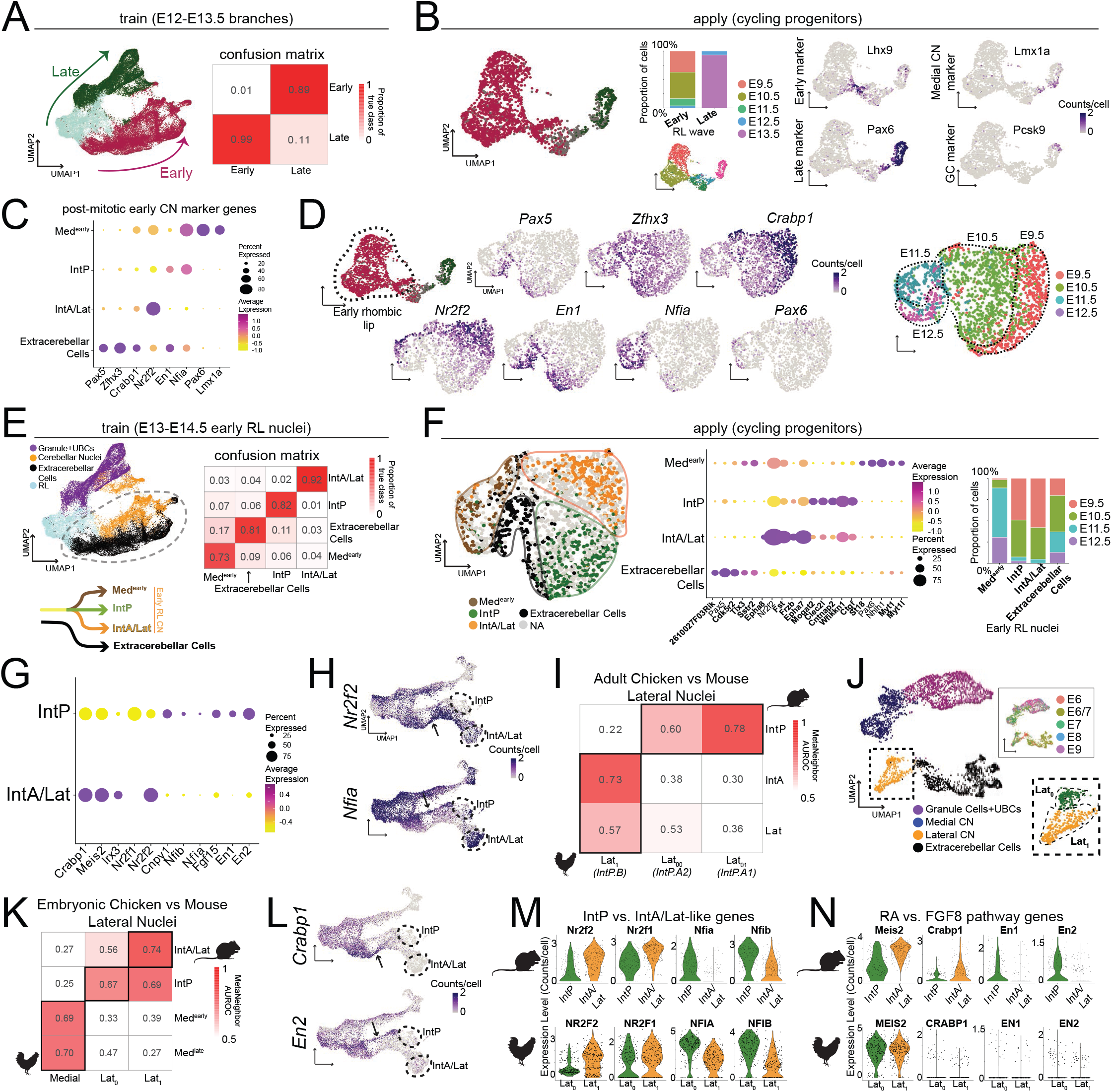
Excitatory cycling progenitors are nucleus-specific with divergent molecular programs in conserved cell types accounting for nuclear differences between mouse and chicken. **(A)** Multinomial logistic regression classifier trained on postmitotic early and late rhombic lip populations (left) at E12.5 and E13.5, with confusion matrix on 50% held-out data demonstrating accurate classification (right). **(B)** Application of the classifier to rhombic lip cycling progenitors reveals high-confidence assignment to early and late populations (left), that correlate with expected developmental timepoints (middle), with corresponding gene expression patterns (right). **(C)** Dot plot of differentially expressed genes within postmitotic early rhombic lip nuclei populations at E13.5. **(D)** UMAP embedding of cycling progenitors confidently assigned to early rhombic lip from **(B)**, colored by expression of selected differentially expressed genes from **(C)** (left) and by timepoint (right). **(E)** Multinomial logistic regression classifier trained on postmitotic nuclei derived from the early rhombic lip at E13.5 and E14.5 (left), with confusion matrix on held-out data (right). **(F)** Application of the classifier to early rhombic lip cycling progenitors from **(B)** shows high-confidence assignments that segregate in UMAP space (left), with differentially expressed genes across putative progenitor cell types (middle, genes not used to train the classifier are in bold), and differential representation across developmental timepoints (right). **(G)** Differentially expressed genes between putative posterior interposed (IntP) and anterior interposed and lateral (IntA/Lat) cerebellar nuclei progenitors. **(H)** Expression of IntA/Lat specific gene *Nr2f2* and IntP specific gene *Nfia* during all developmental stages in postmitotic excitatory cerebellar nucleus UMAP embedding. **(I)** MetaNeighbor AUROC heatmap between adult chicken lateral cerebellar nucleus cell types and adult mouse interposed and lateral cell types, demonstrating that chicken cerebellar nuclei already contain cell types homologous to mouse IntP and IntA/Lat. Chicken cell type names and mouse nucleus labels from ^3^ are provided in parenthesis. **(J)** UMAP embedding of RL-derived excitatory cell types in chicken, with Lat0 and Lat1 division shown in the inset. **(K)** MetaNeighbor AUROC heatmap between cerebellar nucleus populations in chicken and mouse at matched developmental timepoints (E6/7 in chicken, E13.5 in mouse). **(L)** Expression of RA pathway gene *Crabp1* and FGF8 pathway gene *En2* across developmental stages in the postmitotic excitatory cerebellar nucleus UMAP embedding. **(M)** Expression of *Nfi* family genes versus *Nr2f* family genes in postmitotic E13.5 mouse embryonic clusters and corresponding postmitotic E6/7 chicken Lat0 and Lat1 clusters. **(N)** Expression of RA pathway genes (*Meis2*, *Crabp1*) and FGF8 pathway genes (*En1*, *En2*) in postmitotic E13.5 mouse IntA/Lat and IntP populations, respectively, but not in corresponding postmitotic E6/7 chicken lateral nucleus clusters.

**Figure 5.**
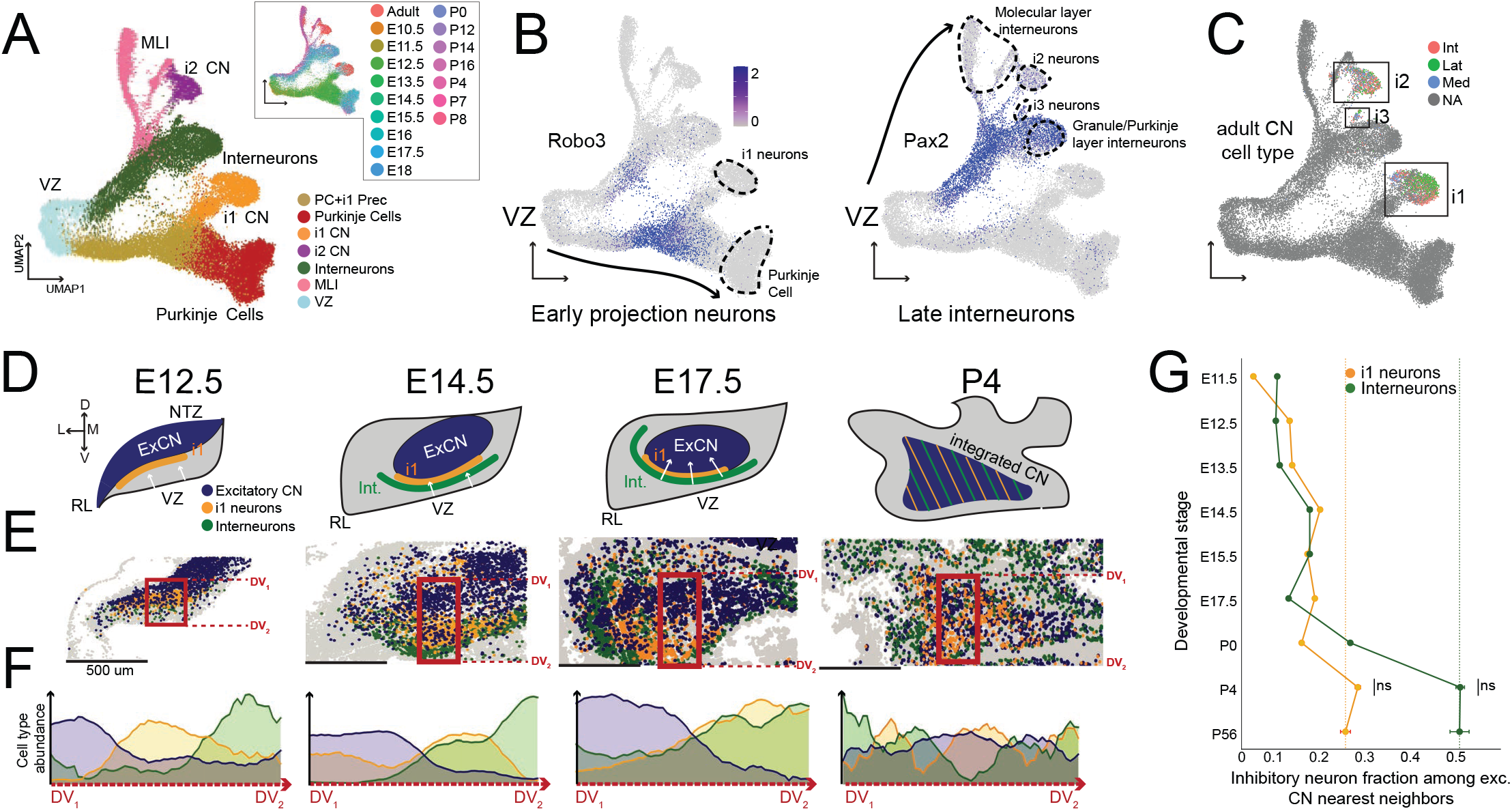
Inhibitory neurons integrate into established nuclear territories. **(A)** UMAP embedding of mouse cerebellar inhibitory neurons from E10.5 to adult, revealing major cell types including Purkinje cells, cerebellar nuclei inhibitory neurons (i1, i2, i3), intermediate Purkinje cell and i1 precursors (PC+i1 Prec), interneurons, molecular layer interneurons (MLI), and postmitotic precursors at the ventricular zone (VZ). **(B)** Two major branches in the cerebellar inhibitory neuron dataset are distinguished by expression of *Robo3* (early projection neurons, left) and *Pax2* (late interneurons, right). **(C)** The same embeddings showing the nucleus identity of adult cells within this plot. Nucleus and cell type labels come from ^3^. **(D)** Schematic summarizing the spatial migration patterns of inhibitory cerebellar nucleus cell types during development. **(E)** Representative BARseq3 slices showing the spatial distribution of excitatory cerebellar nucleus neurons (blue), i1 neurons (orange), and interneurons (green) at representative timepoints E12.5, E14.5, E17.5, and P4. **(F)** Quantification of cell type abundances along the demarcated dorsoventral axis for populations shown in **(E)**. **(G)** Quantification of spatial integration of inhibitory and excitatory cerebellar nucleus neurons, showing the fraction of neighbors to excitatory cerebellar nucleus neurons that are i1 neurons (orange) or interneurons (green) across all sampled developmental stages. Dashed line represents the adult neighborhood state. Wilcoxon rank-sum test shows significantly different neighborhoods until P4. N=2-4 sections per timepoint.

During development, the RL generates extracerebellar-fated neurons, excitatory cerebellar nuclei neurons, unipolar brush cells, and granule cells^18,36^. Around E12.5, it undergoes a major transcriptomic transition that roughly coincides with the onset of granule cell production, dividing the RL and cells produced from it into “early” and “late” waves^37–39^. To understand when and how excitatory neurons fated for the different cerebellar nuclei differ in their gene expression, and how they map onto the different waves of cells produced from the RL, we analyzed mouse scRNAseq data of *Atoh1*-lineage flow-sorted RL-derived neurons spanning E10.5–E16.5^40^ (**Fig. 2A-C; Extended Data Fig. 2A,B; Supplementary Fig. 2A,B**). This dataset contained 52,648 postmitotic neurons from the upper RL, including the expected extracerebellar-fated and cerebellar nuclei neurons, as well as unipolar brush and granule cells. Early and late RL-derived neurons separate into two major branches containing extracerebellar neurons and granule cells, respectively, with the cerebellar nuclei neurons contributing to both (**Fig. 2A**). The early and late RL waves are marked by *Lhx9* and *Pax6* respectively (**Fig. 2B**), along with other differentially expressed genes (**Fig. 2C; Extended Data Fig. 3A**).

To resolve individual cerebellar nuclei and extend our analysis time window, we integrated cerebellar nuclei neurons from this dataset with those of a newly generated E17.5 snRNAseq dataset and a published adult cerebellar nuclei dataset^3^ (**Fig. 2D**; **Extended Data Fig. 3C,E,F; Supplementary Fig. 2C,D,L**). This analysis revealed two major branches of cerebellar nuclei neurons radiating out from the RL, corresponding to the early and late RL waves based on transfer of labels from the whole excitatory cerebellum scRNAseq dataset and expression of marker genes *Pax6* and *Lhx9* (**Fig. 2E,F**). The late RL branch terminates only in the adult medial nucleus, the phylogenetically oldest of the cerebellar nuclei. In contrast, the early RL branch gives rise to all other cerebellar nuclei, including the mammalian-specific and likely phylogenetically youngest IntA and lateral nuclei. This branch splits initially into two trajectories that terminate in IntP and a combination of IntA and the lateral nucleus (IntA/Lat). Later during development, the IntA and lateral nucleus then resolve into distinct identities, mirroring the transcriptomic similarities of the adult nuclei^3^ and their potential evolutionary relationships^7,8^. Interestingly, we could not transcriptionally distinguish between the two excitatory subtypes (Class A and B) that populate each adult nucleus^3^ at any developmental timepoint sampled, suggesting that they emerge within each cerebellar nuclei trajectory after E17.5.

To evaluate whether the same transcriptomic signatures exist in species with a different number of cerebellar nuclei, we analyzed our developmental scRNAseq dataset from chicken, a species which contains only two cerebellar nuclei, and found clusters based on matched orthologous marker genes across species. This analysis revealed a late RL-derived, PAX6+ medial nucleus and an early RL-derived, LHX9+ chicken lateral nucleus, which is thought to be homologous to the mammalian IntP^3^ (**Extended Data Figs. 2E,F; 3B**).

Across amniotes, the late RL wave is thus conserved and only produces the medial nucleus. In contrast, the early RL wave produces more nuclei in mice than chickens, accounting for the different number of nuclei in the adult animals. Taken together, this suggests that the evolutionary addition of the mammalian specific nuclei arose from the expansion of the early RL.

### Nucleus-specific gene expression precedes nuclear territory formation by excitatory neurons

Two scenarios could explain how distinct cerebellar nuclei are generated by the RL during development. First, excitatory neurons destined to form new nuclei could be transcriptomically different from birth in the RL. Alternatively, transcriptomically homogeneous neurons could acquire nuclear identities only after they reach the nuclear transitory zone, for example in response to local external stimuli. The two scenarios point to different evolutionary mechanisms of region formation^8^. To distinguish these possibilities, we analyzed excitatory neurons in our BARseq3 spatial transcriptomics data from E11.5, when the nuclear transitory zone first starts to form, to E17.5, when all major nuclei are transcriptomically distinct. We assigned cell type identities based on semi-supervised clustering at each time point, correlation of clusters across consecutive timepoints, and marker gene expression (**Fig. 3A,B; Extended Data Figs. 4, 5**; **Supplementary Figs. 3, 4;** *Methods*). By projecting all cells onto the mediolateral axis at each timepoint (**Fig. 3C,D**), we quantified the progressive spatial segregation of cerebellar nuclei populations. At each timepoint, we further matched the spatial gene expression with clusters from our scRNAseq dataset (**Fig. 3E**; **Extended Data Fig. 5**), with the two independent datasets corroborating and extending each other.

At E11.5 and E12.5, the nuclear transitory zone is dominated by extracerebellar-fated neurons (*Zfhx3*+, black), with precursors to early RL-derived cerebellar nuclei (yellow) just exiting the RL (**Extended Data Fig. 4A; Supplementary Fig. 3**). At E13, cells expressing IntP (*Olig2*+, green) and IntA/lateral nucleus (*Ntng1*+, orange) markers become transcriptomically distinct at the nuclear transitory zone based on the genes profiled in our BARseq3 dataset. At E13.5, the medial nucleus (*Lmx1a*+/*Meis2*+, blue) arising from the late RL population exits the RL and migrates in the subpial stream (**Supplementary Figs. 3; 4**). At this stage, the IntA and lateral nuclei-fated cells are still transcriptomically indistinguishable based on both scRNAseq full transcriptome and profiled BARseq3 genes. At E14.5, a dramatic spatial reorganization occurs in the cerebellar anlage, with interposed and lateral nuclei neurons moving to acquire a lateral position and medial nucleus neurons migrating through the subpial stream, over the nuclear transitory zone to acquire a more medial position. This rearrangement results in the cerebellar nuclei populations acquiring their characteristic mediolateral positions seen in the adult. The transcriptomic distinction between IntA and lateral nuclei based on scRNAseq full transcriptome and profiled BARseq3 genes emerges last around E15.5, by which time all cerebellar nuclei display distinct spatial positions and transcriptomic profiles matching their adult identities.

Surprisingly, in addition to the canonical nuclei described above, we also identified an extra early RL-derived cerebellar nucleus domain in our spatial data (**Fig. 3D**, brown). This population first appears in the medial nuclear transitory zone at E13 with features of both lateral and medial nucleus identity: it expresses lateral nucleus markers including *Irx3* and *Pou3f2*, as well as the canonical medial marker *Lmx1a* (**Extended Data Figs. 4; Supplementary Figs. 4**), and ultimately settles in space with the canonical medial nucleus. Guided by this discovery, we identified this population in our scRNAseq data and found that this new medial population first shares gene expression with the other early RL nuclei and then starts expressing signatures of the late RL medial nucleus (**Supplementary Fig. 5**). Gene ontology analysis further revealed significant differences in axon-guidance programs between these two medial populations (**Extended Data Figs. 3D; Supplementary Fig. 5B**). We will refer to this new population as Med^early^ and the canonical late RL medial nucleus population as Med^late^. Importantly, based on MetaNeighbor analysis of clusters at matched developmental timepoints across mouse and chicken, we identified distinct populations of medial cells in developing chickens that correspond better with either Med^late^ and Med^early^, indicating that the Med^early^ population is conserved across amniotes (**Supplementary Fig. 5C**). We further investigate the evolutionary and functional implications of this previously unknown population as described in **Fig. 6**. However, its presence shows that all adult mouse cerebellar nuclei, including the medial nucleus, receive contributions from the evolutionarily dynamic early RL.

**Figure 6.**
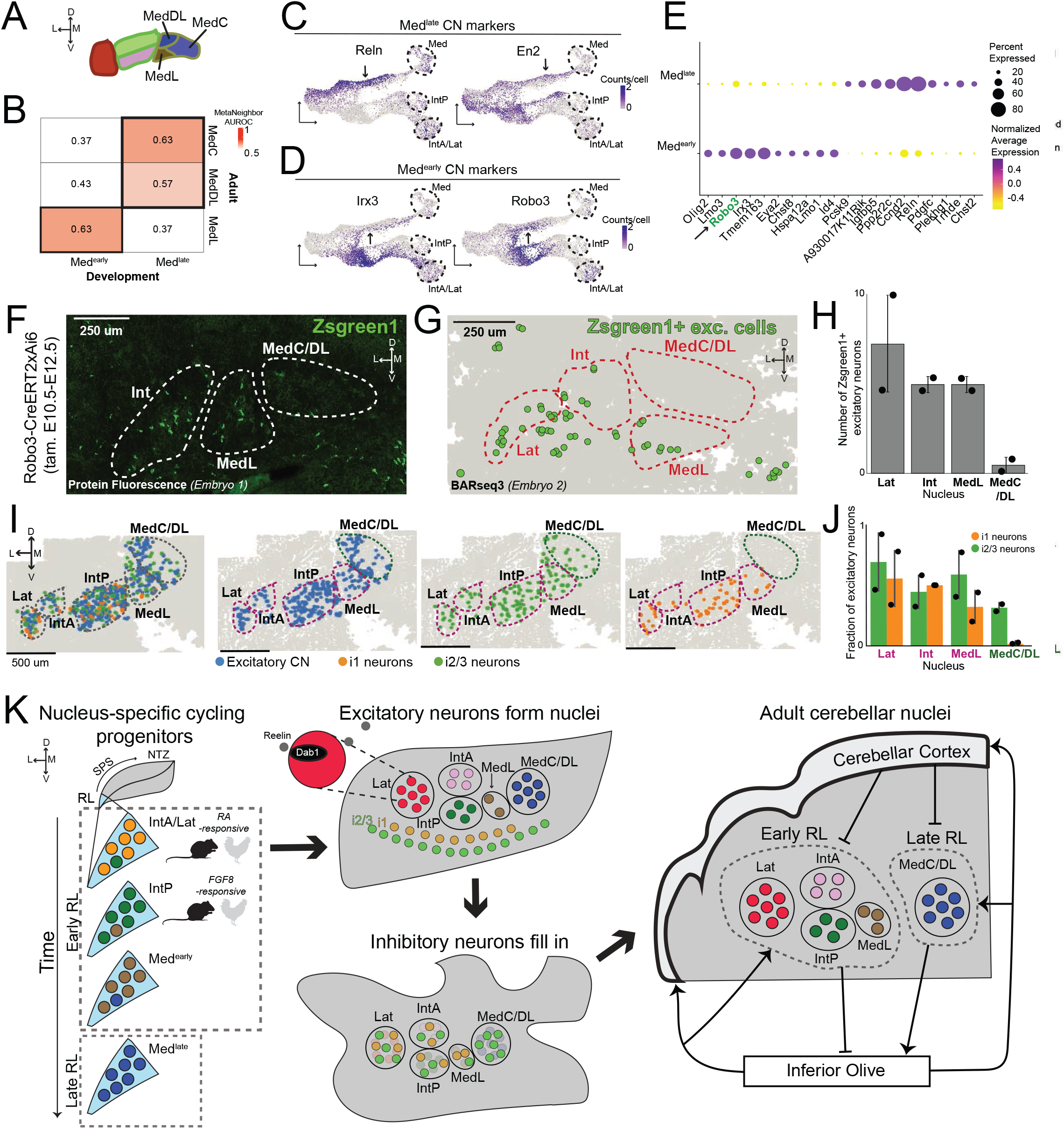
The late rhombic lip gives rise to the higher-order parts of the medial nucleus as part of a non-canonical olivocerebellar circuit. **(A)** Schematic of adult mouse medial cerebellar nucleus subdivisions, including the central (MedC), dorsolateral (MedDL), and rostrolateral (MedL) subnuclei. **(B)** MetaNeighbor AUROC heatmap showing transcriptomic correspondence between Med^early^ and Med^late^ populations at E13.5 and adult medial subnuclei defined by ^3^. **(C,D)** Expression of Med^late^ **(C)** and Med^early^ **(D)** marker genes in UMAP space. Med, whole medial nucleus. **(E)** Dotplot showing differentially expressed genes between Med^early^ and Med^late^ in the E12.5 and E13.5 scRNAseq dataset. **(F)** Representative image of a coronal section of the cerebellar nuclei in *Robo3*-CreERT2×Ai6 mouse embryo 1 at E18.5 showing preferential Zsgreen1 labeling of MedL [in addition to interposed (Int, shown) and lateral (Lat, not shown) nuclei], with minimal labeling of MedC or MedDL. N=2 embryos. **(G,H)** Representative coronal BARseq3 section of the cerebellar nuclei in *Robo3*-CreERT2×Ai6 mouse embryo 2 at E18.5. (**G**) Location of *ZsGreen1*+ excitatory neurons. (**H**) Quantification of *ZsGreen1*+ excitatory neurons shows strong labeling in lateral (Lat), interposed (Int), and MedL nuclei, but minimal labeling in MedC/DL. N=2 sections. **(I)** Representative BARseq3 section of the adult cerebellar nuclei, showing all excitatory and inhibitory neurons (left), and separately only excitatory neurons (middle left), i2/3 neurons (middle right), and i1 neurons (right). Nuclei boundaries are colored by assignment to early or late RL origin. Lat, lateral nucleus. **(J)** Quantification of i1 projection and i2/3 local inhibitory neurons in the adult cerebellar nuclei. i2/3 neurons are present in all nuclei, whereas i1 neurons are virtually absent from MedC/DL. N=2 sections. **(K)** Model of cerebellar nucleus development and evolution supported by the data presented here.

Taken together, our spatial analysis shows that cerebellar nuclei excitatory neuron identities reveal themselves before they reach the nuclear transitory zone: Cells that will ultimately end up in the same cerebellar nucleus are spatially clustered and migrate together, with nucleus-specific transcriptomic identities preceding the final spatial localization at the nuclear transitory zone. By E15.5, excitatory neurons have established all cerebellar nuclear territories. Cerebellar nuclei identities are thus likely not set by a single patterning event at the nuclear transitory zone but must arise earlier. Together with the finding that the early RL gives rise to phylogenetically younger nuclei (**Fig. 2**), these results support a hypothesis where evolutionary innovation leading to new region formation by excitatory neurons occurs in early RL progenitors.

### Early rhombic lip progenitors already exhibit transcriptomic signatures of individual nuclei

To test this hypothesis, we asked whether excitatory cerebellar nuclei identity is specified within cycling RL progenitors or emerges only after cell-cycle exit. We initially trained a multinomial logistic regression classifier on E12.5 and E13.5 scRNAseq data from postmitotic neurons to distinguish early and late RL identities, and applied it to a scRNAseq dataset of cycling RL progenitors, identified as E9.5 to E13.5 *Atoh1*+ cells that express *Mki67* and do not express *Hox* genes^40^. As expected from the large transcriptomic changes in the RL between its early and late states^37–39^, this classifier confidently assigned many cycling cells to early or late cerebellar nuclei programs (**Fig. 4A,B; Extended Data Fig. 6A-C,E-F**). Early RL progenitors predominated from E9.5 to E11.5, whereas late RL progenitors were enriched from E12.5 to E13.5 (**Fig. 4B**, *middle*), matching the temporal progression observed among postmitotic cerebellar nuclei populations. Many assigned progenitors expressed the expected wave-associated markers, including *Lhx9* and *Pax6* (**Fig. 4B**, *right*).

We next asked whether nuclear diversity is already evident in cycling progenitors that were confidently assigned to the early RL. First, we performed unsupervised clustering of RL progenitors, which showed substantial transcriptional heterogeneity within this population (**Extended Data Fig. 6G,H**). Next, we wanted to evaluate whether any of this heterogeneity corresponded to cerebellar nuclei identities. Genes differentially expressed among Med^early^, IntP, and IntA/Lat postmitotic cerebellar nuclei populations at E13.5 (**Fig. 4C**) also varied across cycling early RL progenitor clusters from E9.5 to E12.5 (**Fig. 4D**). To explicitly assign progenitors to future cerebellar nuclei identities, we then trained a second classifier on E13.5 and E14.5 postmitotic early RL-derived neurons and applied it to cycling early RL progenitors (**Fig. 4E**). This classifier confidently assigned many progenitors to Med^early^, IntP, IntA/Lat, and extracerebellar cell identities prior to cell-cycle exit (**Fig. 4F**, *left*; **Extended Data Fig. 6A,B,D-F**), In contrast, IntA and lateral nucleus identities could not yet be reliably distinguished (see *Methods*), consistent with the late transcriptomic separation of these two cell populations (**Figs. 2; 3**). Importantly, putative nuclei-specific progenitor populations differentially expressed many genes, including those not used to train the classifier, indicating these are biologically distinct progenitors rather than noisy classifier predictions (**Fig. 4F**, *middle*).

The predicted progenitor identities were differentially represented across developmental time in overlapping windows (**Fig. 4F**, *right*). Putative IntA/Lat progenitors were enriched at the earliest stages, followed sequentially by putative IntP and Med^early^ progenitors. Extracerebellar fated progenitors, in contrast, spanned E9.5-E12.5, consistent with prior models of temporal birth dating from RL-derived populations^18,39,41,42^. Taken together with our spatial analysis (**Fig. 3**), this suggests that excitatory cerebellar nuclei neurons emerge from the rhombic lip in an order consistent with their lateral to medial positioning in the adult: IntA/Lat first, IntP second, Med^early^ third, and Med^late^ last. Overall, early and late RL progenitors exhibit temporally ordered transcriptional heterogeneity associated with future cerebellar nuclei identities. Thus, the evolution of new cerebellar nuclei was likely mediated by expansion and diversification of distinct early RL progenitor pools.

### Cell type generation and nucleus formation are independent events

To investigate the transcriptional programs that may have supported the formation of new cerebellar nuclei during evolution, we focused on the development of the excitatory neurons of the putatively mammalian-specific IntA/Lat cerebellar nuclei compared to their conserved sister nucleus IntP. IntP and IntA/Lat progenitors and postmitotic trajectories are defined by key transcription factors, including *Nfia/b* for IntP and *Nr2f1/2* for IntA/Lat (**Fig. 4G,H**). These transcription factors have been implicated in neuronal specification in other brain regions, including in the retina, where they act as terminal transcription factors with *Nr2f2* marking early-born and *Nfia* marking later-born cell types^43–46^. Interestingly, this temporal progression mirrors the emergence of IntA/Lat progenitors before IntP progenitors in the RL (**Fig. 4F**, *right*).

Chickens have only two cytoarchitectonic cerebellar nuclei, and in previous work, we showed that the excitatory pseudobulk transcriptome of the adult chicken lateral nucleus most closely matches to that of the mammalian IntP, suggesting homology between these two regions and de novo emergence of the mammalian IntA/Lat^3^. However, finer comparison shows that the different excitatory cell types that make up the chicken lateral nucleus transcriptomically match best to either adult mouse IntP or IntA/Lat [^3^ and **Fig. 4I**]. Similarly, in our chicken developmental dataset, we detected two early RL-derived chicken cerebellar nuclei cell types (Lat_0_ and Lat_1_) that matched to mouse IntP or mouse IntA/Lat, respectively, (**Fig. 4J,K)** driven by *Nfia/b* and *Nr2f1/2* gene expression programs (**Fig. 4M**). The cell types forming IntP and IntA/Lat are therefore conserved between chickens and mice, even though the distinct nuclei are not. Accordingly, cell type generation and the physical sorting of cell types into new nuclei are likely independent events in evolution.

### Mammalian-specific co-option of FGF8 and RA signaling in conserved cell types

To investigate how conserved cell types may be sorted into different cerebellar nuclei, we examined differentially expressed genes between the putative IntP and IntA/Lat progenitors in mice. Interestingly, we detected enrichment of genes linked to fibroblast growth factor-8 (FGF8) and retinoic acid (RA) signaling pathways in IntP and IntA/Lat progenitors, respectively (**Fig. 4G**). FGF8 and RA are well-established mutually antagonistic signaling pathways that regulate boundary formation in many different developmental contexts, including the development of the retina, limb bud, and general body axis segmentation^47–50^. IntP progenitors preferentially expressed FGF8-associated genes, including *En1*, *En2*, *Fgf15*, and *Cnpy1*, whereas IntA/Lat-associated progenitors preferentially expressed RA-associated genes, including *Nr2f2*, *Crabp1*, and *Meis2*. Many of these transcriptional differences persisted through later maturation in both scRNAseq and spatial datasets (**Fig. 4L; Extended Data Fig. 7A**).

We analyzed published mouse snATAC-seq datasets spanning E12.5 to E14.5^28^ to assess whether these differences in RA-FGF8 gene expression programs were associated with chromatin accessibility differences (**Extended Data Fig. 7B-H**). Indeed, we found that the RARE motif (target of RA pathway) was significantly more enriched in the IntA/Lat population compared to IntP, with support from footprinting analysis (**Extended Data Fig. 7J-L**). Given that RA is primarily synthesized in the meninges surrounding the developing cerebellum, and FGF8 is produced at the isthmus, and considering the small size of the RL, spatial gradients of these diffusible signals are unlikely to explain the observed differences^51–54^. Overall, differences between IntP and IntA/Lat populations may thus reflect distinct RA-response competencies among early RL-derived progenitors rather than the differential exposure to spatial gradients of diffusible FGF8 and RA.

Strikingly, despite the presence of cell types homologous to IntP and IntA/Lat in the chicken lateral nucleus, we did not detect the RA- and FGF8-associated transcriptional programs in these cells (**Fig. 4N**). This difference is unlikely to be caused by technical factors (**Extended Data Fig. 8A-E**), and is consistent with a previous study showing that RA-pathway elements (i.e. CRABP1) are absent from the cycling RL progenitors of the chicken cerebellum, but are present in the mouse RL^54^. Evolutionary co-option of RA- and FGF8-associated programs in mammals in conserved cell types may thus contribute to the spatial separation of nuclear populations in the mammalian lineage and the formation of physically separated IntP and IntA/Lat cerebellar nuclei. Taken together, these findings suggest that evolutionary brain region duplication can occur through a process where conserved sister cell types from an ancestral region diverge and undergo a "regionalization event" in which they are sorted into distinct cytoarchitectonic structures.

### The divergence of the excitatory cells of the anterior interposed and lateral nuclei

Complementing this early nucleus division, we sought to understand the factors driving the late separation of IntA and the lateral nucleus in mice. Analysis of differential gene expression between these populations at their divergence at E15.5 and E17.5 reveal potential differences in Reelin signaling (**Extended Data Fig. 8F**). The lateral nucleus expresses *Dab1*, which is activated when Reelin binds to extracellular receptors, while IntA neurons do not. At these timepoints, granule cells, which are the primary *Reelin*-expressing cell type, become abundant and surround the cerebellar nuclei^55–57^. Because Reelin-Dab1 signaling regulates neuronal migration ^58,59^, this difference could contribute to the spatial segregation of these populations. Supporting this suggestion, *reeler* mice homozygous for a *Reelin* mutation show improper migration and loss of demarcation of only the lateral nucleus^60–62^.

### Inhibitory neurons fill in after the excitatory cells

Above, we showed that excitatory neurons establish nuclei architecture by E15.5, making them a strong candidate for causing evolutionary changes in cerebellar nuclei number. However, over half of all neurons in the adult cerebellar nuclei are inhibitory neurons, and these cells could also contribute to region formation alongside the excitatory neurons. In adults, each inhibitory neuron class is transcriptomically invariant across nuclei, but this does not preclude nucleus-specific gene expression trajectories and a function in defining cerebellar nuclei territories during development^63^. To test whether inhibitory neurons show nucleus-defining roles during development, and potentially evolution, we examined their gene expression and migration in the developing cerebellum.

Inhibitory cerebellar nuclei neurons are born from the *Ptf1a*+ ventricular zone of dorsal rhombomere 1^21–23^. This same developmental niche gives rise to Purkinje cells and cerebellar cortical interneurons over a protracted time course: Inferior olive-projecting i1 cerebellar nuclei neurons and Purkinje cells are born around E10-12.5 in mice, and local i2 and i3 cerebellar nuclei and cortical interneurons are born later until birth^35,64,65^. To understand the developmental trajectories of these populations, we integrated the inhibitory neurons of five published scRNAseq datasets^3,27–30^ with E17.5 snRNAseq data generated as part of this study to obtain a comprehensive dataset of 31,279 inhibitory neurons covering E10.5 to adulthood (**Fig. 5A; Supplementary Fig. 2E,F,L; Extended Data Figs. 2C,D; 9**).

The embedding revealed two developmental branches, corresponding to the “early” and “late” ventricular zone-derived inhibitory populations (**Fig. 5B**)^35,66^. The early branch, emerging at E12.5, produces the inferior olive-projecting i1 neurons of the cerebellar nuclei and the Purkinje cells of the cerebellar cortex. The late branch, emerging at E17.5, produces later-born interneuron populations of the cerebellar nuclei and the cerebellar cortex, including the glycinergic i2 and i3 cerebellar nuclei neurons and molecular layer interneurons (**Fig. 5A,C**). Similar to the excitatory cerebellar nuclei neurons, therefore, inhibitory cerebellar nuclei neurons are also born across early and late states of their germinal zone. However, unlike excitatory cerebellar nuclei neurons, their developmental trajectories reflect inhibitory neuron subtype rather than cerebellar nucleus identity.

To determine how inhibitory neurons spatially interact with developing excitatory cerebellar nuclei neurons, we analyzed BARseq3 data across all developmental timepoints to account for their protracted development. Throughout embryonic development, inhibitory cerebellar nuclei neurons remained largely segregated from excitatory cerebellar nuclei populations, with i1 neurons occupying the territory directly ventral to the excitatory nuclear transitory zone, as previously described^65^, and interneurons (including i2/3) occupying the space ventral to the i1 neurons (**Fig. 5D-F***, E12.5, E14.5, E17.5;* **Extended Data Fig. 9**). At early postnatal timepoints, after the excitatory neurons have set up nuclear territories, both i1 and interneurons began integrating into these cerebellar nuclei territories (**Fig. 5D-F***, P4;* **Extended Data Fig. 9**). To quantify inhibitory neuron integration into excitatory cerebellar nuclei territories, we calculated the ratio of inhibitory to excitatory cerebellar nuclei neurons within each excitatory neuron’s twenty nearest neighbors across all cerebellar nuclei. We then asked when these local neighborhood compositions became statistically indistinguishable from the adult pattern. This quantification demonstrated progressive integration of both i1 and interneuron populations, starting at late embryonic timepoints and reaching distributions comparable to the adult state by P4 (**Fig. 5G; Extended Data Fig. 9**).

Importantly, although inhibitory neurons can be transcriptionally resolved into i1, i2, and i3 populations prior to integration based on the scRNAseq embedding (**Fig. 5C**), we found no evidence of transcriptomic heterogeneity indicative of their assignment to a specific cerebellar nucleus. Instead, excitatory neurons establish the molecular and spatial scaffold of individual cerebellar nuclei first, with inhibitory populations integrating into these pre-established structures during early postnatal development (**Fig. 5D-G**). The evolution of new cerebellar nuclei is therefore driven primarily by elaborations of the early RL, giving rise to specialized excitatory neurons that form new cerebellar nuclei territories which then get “filled in” by nucleus-invariant inhibitory neurons.

### Dual origin of the medial nucleus explains its divergent functions

The function and evolutionary status of the mammalian medial nucleus remain to be understood. Traditionally, the medial nucleus has been viewed as an ancient structure with vestibular and brainstem functions, yet, clinical and basic research work in mice and humans conclusively demonstrates its function in higher cognitive domains, including working memory, motivation and fear behaviors^11–16^. Our work here reveals that the adult medial nucleus is formed by two distinct developmental populations: the canonical late RL-derived Med^late^ and the newly discovered early RL derived Med^early^ (**Fig. 3D**). We hypothesize that this dual developmental and likely evolutionary origin of the medial nucleus is at the root of its simultaneous “old” vestibular/brainstem and “new” cognitive functions. To test this idea, we sought to identify which part of the adult medial nucleus is formed by Med^early^ and Med^late^. The adult medial nucleus can be divided into three subnuclei, the MedDL (dorsolateral protuberance of the medial nucleus), MedL (rostrolateral division of medial nucleus), and MedC (central medial nucleus) (**Fig. 6A**)^3^. While MedC and MedDL have been associated with the higher-order functions of the medial nucleus based on their projection patterns, the MedL projects to the brainstem^13,67^, likely carrying out more basal functions^13^.

Transcriptomically, E13.5/14.5 Med^early^ showed strongest correspondence with adult MedL, whereas Med^late^ corresponded to MedC and MedDL (**Fig. 6B**). To directly test where Med^early^ neurons settle in the adult, we performed genetic lineage tracing. Like the other early RL derived cerebellar nuclei, Med^early^ expresses *Robo3* during early development (E10.5 to E12.5), while Med^late^ does not (**Fig. 6D,E**). In addition to the cerebellar nuclei excitatory neurons, *Robo3* is expressed in various inhibitory neurons during this time window (**Fig. 5B**). To permanently label Med^early^ neurons, we crossed *Robo3*-CreERT2 to Ai6 *Zsgreen1* Cre-reporter mice and administered tamoxifen to activate Cre from E10.5 to E12.5. At E18.5, when cerebellar nuclei boundaries are cytoarchitectonically apparent, *ZsGreen1*+ cells in the medial nucleus were primarily confined to the MedL subnucleus and were absent from MedC/DL (**Fig. 6F; Supplementary Fig. 6**), suggesting that MedL but not the other medial subnuclei are early RL-derived. To exclude the possibility that this result was due to labeling of inhibitory neurons that already integrated into the nucleus, we performed BARseq3 on similar samples using a targeted 31-gene panel containing cerebellar cell type markers. Within the medial nucleus, *ZsGreen1*+ excitatory neurons were present in the MedL subnucleus, while the MedC/DL contained almost none (**Fig. 6G,H; Extended Data Fig. 10A-C**), supporting our previous conclusion. Complementary support for the Med^early^ population becoming MedL comes from previous *SepW1*-Cre; *En1*/*En2* knockout studies^33,68^, which we predict should specifically disrupt Med^late^ based on its gene expression and indeed show loss of MedC/DL while leaving MedL largely intact (**Fig. 6C; Extended Data Fig. 10E,F**).

Early RL-derived Med^early^ thus gives rise to the brainstem projecting MedL, whereas late RL-derived Med^late^ gives rise to the thalamus projecting MedC/DL. Within the evolutionarily expanding early RL-derived cerebellar nuclei, the traditional notion holds; the ancient medial nucleus performs basal functions and newer, more lateral nuclei perform higher-order functions. By contrast, the late RL gives rise to a part of the medial nucleus outside of this scheme to endow the mammalian medial nucleus with its higher-order functions.

### Only early rhombic lip derived cerebellar nuclei are part of the canonical olivocerebellar loop

The inhibitory neurons of the cerebellar nuclei are critical components of the closed olivocerebellar loop of the cerebellum: i1 cerebellar nuclei neurons project to the inferior olive, which in turn project to Purkinje cells in the cerebellar cortex, which then complete the loop by projecting to the cerebellar nuclei^69,70^. This loop is thought to be central to the cerebellar computation, particularly in its involvement in predictive error to refine movements^71^.

Given the status of the late RL-derived cerebellar nuclei as falling out of the overall pattern set up by the early RL derived nuclei, we wondered if these different cerebellar nuclei are equally populated by inhibitory neurons. When we quantified the abundance of i2/3 neurons per subnucleus in our adult BARseq3 dataset, we found that all cerebellar nuclei are populated by these interneurons (**Fig. 6I,J**). Strikingly, however, the inferior olive-projecting i1 neurons are abundant in the early RL-derived lateral, interposed, and MedL but are specifically absent from the late RL-formed MedC and MedDL (**Fig. 6I,J; Extended Data Fig. 10G**). Similarly, i1 neurons labeled by tamoxifen administration from E10.5 to E12.5 in *Robo3-CreERT2xAi6* mice are located only proximal to parts of the cerebellar nuclei that contain early RL-derived excitatory neurons before integration at E18.5 (**Extended Data Fig. 10B,C**). These findings are consistent with previous reports that *Sox14*+ i1 neurons are concentrated in the rostroventral medial compartment of the medial nucleus^65,72^, and that the inferior olive-projections from large parts of the medial nucleus are uniquely excitatory, rather than inhibitory^72^. The olivocerebellar loop, with inhibitory cerebellar nuclei to inferior olive projections, and hence the canonical cerebellar circuit, is therefore only present in the cerebellar nuclei formed by the early RL. In contrast, the late RL-derived, higher-order, parts of the medial nucleus form a different cerebellar circuit altogether.

## DISCUSSION

Here we provide a developmental framework for understanding how new cerebellar nuclei, each containing a conserved cell type-set in the adult animal, are generated across vertebrate evolution (**Fig. 6K**). We show that evolutionary region duplication is driven by excitatory neurons from the early RL, and that new cerebellar nuclei are likely formed when sister cell types diverge and become spatially segregated into distinct brain regions. Later during development, inhibitory neurons fill in the cerebellar nuclei territories set up by the excitatory cells to complete the cell type set and form the canonical cerebellar circuit. Intriguingly, a special case of this rule is found in the medial cerebellar nucleus, in which the subset of subnuclei that are responsible for this nucleus’ non-canonical higher cognitive functions is set up by late RL derived excitatory cells and miss a key inhibitory cell type, thus forming an alternative cerebellar circuit.

### Excitatory neurons lead the way

Generation of new cerebellar nuclei requires the assembly of excitatory and inhibitory neurons born from distinct developmental niches. We found that this process is strictly hierarchical; excitatory cells physically establish all cerebellar nuclei territories by E15.5, and their specific regional identities are largely discernible in the cycling progenitor cells of the RL. In contrast, inhibitory neurons integrate into the established cerebellar nuclei territories only at early postnatal timepoints, without any apparent cerebellar nuclei specific gene expression signatures. These findings align with previous work showing that the adult excitatory cell types are nucleus-specific, whereas adult inhibitory cell types are nucleus-invariant^3^. Cerebellar nuclei identity, generation, and evolution thus appear to be dictated by excitatory cells.

This excitatory-neuron–driven logic is not unique to the cerebellar nuclei. In the cerebral cortex, excitatory neurons establish transcriptomically and cytoarchitecturally defined layers before inhibitory neurons populate them^73^. Excitatory neurons also diversify across evolution, forming new (sub)layers and changing projection patterns^74–76^ within mammalian cortex, or establishing entirely different pallial architectures, for example in birds vs. mammals^77,78^. In contrast, pallial inhibitory neurons remain remarkably conserved^78–81^. Interestingly, these features are not tied to the physical location where these neurons are generated relative to their final position or their migration route. While cerebral cortical excitatory cells migrate radially and cortical inhibitory cells migrate tangentially, in the cerebellar nuclei excitatory cells migrate tangentially and inhibitory neurons radially. Importantly, the excitatory-first pattern is also not a “projection-neuron-first” pattern. Both cerebral cortical and cerebellar nuclei excitatory neurons are generally projection neurons. However, whereas cortical inhibitory neurons are almost always local interneurons, the i1 inhibitory neurons of the cerebellar nuclei are projection neurons and nevertheless follow the lead of the excitatory neurons. It should be noted, however, that i1 neurons project only within the olivocerebellar system, while excitatory neurons project more broadly. Overall, excitatory neurons thus appear to be the primary substrate for evolutionary modification across much of the vertebrate brain.

### New cell types before new regions

Given this excitatory-cell–driven model, the existence of nucleus-specific progenitors in the early RL and their sister cell type status to each other^3,7^, new cerebellar nuclei may be formed (i) de novo by generating a novel progenitor specific to a new nucleus or (ii) via a gradual path where the progenitor pool first diversifies to generate new sister cell types within the same cerebellar nucleus, and the sister cell types then diverge to sort into new spatially separate regions. Our analysis of IntP and IntA/Lat development across mouse and chicken showing a mammalian specific co-option of RA/FGF8 signaling in conserved cell types support the latter, two-step model. However, when exactly the new sister cell types emerged during evolution of the cerebellar nuclei or if they were inherited from other regions remains unknown.

This two-step model provides an important contribution to the duplication and divergence model of the cerebellar nuclei. Brain regions composed of repeated cell type units that appear as “duplicates” of each other in the adult brain are not formed in a simple, instantaneous duplication that precedes functional divergence, breaking the analogy of region and gene duplication. Instead, these duplications can have their origins in ancestral brain regions that contain overall a similar set of cell types, but that also already contain “seed” cell types that further diverge and are ultimately spatially sorted to produce new brain regions. The functional role of these region precursors remains to ascertained, but their existence suggests that the evolution of novel brain functions may precede the evolution of new regions.

### Only early rhombic lip cerebellar nuclei are part of the canonical cerebellar circuit

A fundamental cerebellar circuit is the olivocerebellar loop, which plays a critical role in prediction error signaling and motor learning^82^. Strikingly, our results demonstrate that this circuit is only formed by (sub)nuclei that are set up by early RL derived excitatory neurons: MedL, IntA, IntP, and the lateral nucleus. The late RL-derived medial subnuclei MedC and MedDL do not recruit i1 neurons for integration into their nuclear territory and thus cannot support the inhibitory cerebellar nuclei to inferior olive projections of the olivocerebellar loop. Consistent with this finding, recent work demonstrates that MedC/DL uniquely sends excitatory but not inhibitory projections to the inferior olive in mice^72^, thus forming a different, sign-inverted, olivocerebellar circuit.

Intriguingly, the distinction between early and late RL-derived subnuclei also resolves a tension in the literature on the function and projection of the mammalian cerebellar nuclei. Much of the classic literature assigned the “ancient” medial nucleus projections to vestibular and brainstem targets and corresponding functions, while the “newer” interposed and lateral nuclei project to thalamic targets and perform higher-order functions. More recent findings challenged this view by highlighting significant thalamic projections and higher-order functions in the medial nucleus^11–16^. Our results now suggest that the classic view does indeed hold true for the early RL-derived cerebellar nuclei that partake in the classic olivocerebellar loop, since MedL has recently been shown to project to the brainstem and not the thalamus^13,67^. In contrast, thalamic projections of the medial nucleus appear to originate from the late RL-derived MedC/DL^13^, which do not form the classic cerebellar circuit. Higher-order, thalamus-projecting portions of the medial nucleus thus are fundamentally different from the rest of the cerebellar nuclei.

## CONCLUSION

Taken together, our findings suggest a simple model for cerebellar evolution: Sister excitatory progenitors of the early RL are generated, diverge, and ultimately sort into new territories. Each territory is then filled in by nucleus invariant inhibitory i1 projection neurons and i2/3 interneurons, establishing a new cerebellar nucleus that partakes in the canonical cerebellar circuit. Based on evidence that excitatory cerebellar nuclei neurons developmentally scale the number of Purkinje and granule cells^33,83^, it is tempting further to speculate that this same mechanism will then “automatically” create the appropriate input to the new cerebellar nuclei. Future work will address this interaction and determine when and how excitatory progenitors diversify to form new cell types.

## METHODS

### Animals

All procedures followed Johns Hopkins University Animal Care and Use Committee (ACUC) protocol MO23M346. We used C57BL/6J (Jackson Laboratory, strain #000664) mice for timed pregnancies in most mouse experiments. To map the early medial population, we crossed Robo3-CreERT2 females (Jackson Laboratory, strain #031215) with Ai6 males (Jackson Laboratory, strain #007906), administered tamoxifen as described below from E10.5 to E12.5, and collected embryos at E18.5. For embryonic staging, we either purchased wildtype timed pregancies from Jackson Laboratory or set up timed pregnancies such that the day of vaginal plug detection counted as E0.5. We included both male and female embryos in experiments. We obtained fertilized white leghorn chicken eggs from commercial sources (University of Connecticut and Michigan State University). We staged chicken embryos according to Hamburger-Hamilton developmental criteria^84^ and collected them at the indicated embryonic stages. We validated findings through technical replicates, including multiple sections per embryo, and through cross-validation between orthogonal methods, including scRNAseq and BARseq3.

### BARseq3 probe design and preparation

#### SNAIL probe design

We designed SNAIL probes as previously described for BARseq3^17^. Briefly, for genes with multiple isoforms, we identified regions conserved across all annotated isoforms and designed probes to target those shared sequences. We used PICKY v2.2^85^ to select unique target sequences and minimize off-target hybridization, and we designed the hybridization arms of each probe pair to be 40–46 nucleotides long. Each SNAIL probe set for a unique gene carried a randomly generated gene-specific ID sequence encoded in the padlock probe. We designed gene ID sequences with a minimum Hamming distance of 2 between genes. We designed four probes for each gene.

#### Probe amplification and preparation

We ordered a custom oligonucleotide pool from Twist Biosciences that contained up to four SNAIL probes per gene and targeted 1,761 genes, including all known transcription factors, genes involved in cellular migration and adhesion, a curated set of neuronal marker genes, and additional targets (**Extended Data Table 1**). We amplified these SNAIL probes using a previously published rolling circle amplification method^17,86^ with modifications to improve final oligo yield. To increase RCA amplification yield, we added 0.16 mg/mL T4 gene 32 protein (NEB M0300L) to each RCA step. We replaced the nicking enzyme and primer sites for Nt.BspQI with those required for Nt.Alw1 digestion and digested the final RCA-amplified samples with 0.3 U of Nt.Alw1 (NEB R0627S) and 0.3 U of Nb.BtsI (NEB R0707S). To increase the yield of fully digested oligo products, we performed three sequential nicking digestions by denaturing the nicking enzymes, rehybridizing the nicking primers, and adding 0.3 U of fresh nicking enzymes for each round. After digestion, we pelleted the samples to remove denatured proteins and then purified the amplified oligos by ethanol precipitation. We rehydrated the samples with RNase-free water, measured final concentrations, and lyophilized the amplified product. We evaluated library concentrations by gel electrophoresis and resuspended the lyophilized oligos to 200 µM in RNase-free water. To increase detection of genes of interest, we added four additional probes per gene for 31 selected genes using a probe panel we ordered from IDT, giving these genes a total of eight probes per gene in transcriptomic experiments (**Extended Data Table 1**). We also used this secondary probe panel by itself to decode cell types in the Robo3-CreERT2 × Ai6 lineage-tracing experiment.

All oligonucleotide sequences are provided in **Extended Data Table 1**.

### BARseq3 tissue and library preparation

#### Fresh-frozen tissue preparation

We performed all BARseq3 experiments on fresh-frozen tissue sections except for those from Robo3-CreERT2 × Ai6 embryos (described below). At the required embryonic stages, we deeply anesthetized pregnant mice or early postnatal mice with Isofluorane, and decapitated them. We dissected embryonic and postnatal brains in cold phosphate-buffered saline (1X PBS) and flash-froze them in optimal cutting temperature (OCT) compound.

We coated glass-bottom plates (MatTek, P24G-1.5-13-F) with Bind-Silane coating solution [0.1% (vol/vol) Bind-Silane (Sigma-Aldrich, 440159-100ML), 80% (vol/vol) ethanol, 2% (vol/vol) glacial acetic acid (Sigma-Aldrich, A6283-100ML) in ultrapure water (Invitrogen, 10977-023)]. We left the coating solution in the wells until it evaporated. We rinsed the wells once with 100% ethanol, then added 1 mg/mL poly-D-lysine (Sigma-Aldrich, A-003-E) to each well. We sealed the plates with Parafilm and incubated them overnight at 4°C. Before cryosectioning, we washed the wells three times with RNase-free water (Invitrogen, 10977-023) and air-dried them.

We cryosectioned samples at 16 µm, melted them directly onto the glass well surface, and refroze them. After cryosectioning, we air-dried the sections at room temperature until they became translucent (∼5 min). We then fixed them in 4% paraformaldehyde (EM-grade PFA; Electron Microscopy Sciences, 15710-S, diluted in 1× PBS; Thermo Fisher Scientific, Gibco 70011044) for 10 min at room temperature and washed them three times with 1× PBS. We then added ice-cold 100% methanol, replaced it with fresh ice-cold 100% methanol, and stored the samples at –80°C for at least 1 hour before warming them to room temperature for library synthesis.

#### Fixed tissue preparation for Robo3-CreERT2 × Ai6 embryos

For Robo3-CreERT2 × Ai6 animals, we dissected E18.5 whole embryos, decapitated them, and fixed the heads in 4% PFA for 48 hours. We then incubated the tissues sequentially in 15% sucrose and 30% sucrose for 24 hours each. After that, we flash-froze the tissues in OCT. We cryosectioned the samples at 20 µm onto Superfrost PLUS slides (Fisher Scientific, 1255015), and installed HybriWell-FL chambers (22 mm × 22 mm × 0.25 mm; Grace Bio-Labs) after sections and slides equilibrated to room temperature.

We treated fixed tissue slices with pepsin [0.2% (wt/vol) pepsin (MilliporeSigma, P7012) in 0.1 M HCl] for 5 min at room temperature and then washed them in 1× PBS. We dehydrated the samples sequentially in 70%, 85%, and 100% ethanol for 5 min each. We then added fresh 100% ethanol to the chambers, incubated the slides at 4°C for 1.5 hours, and performed library synthesis.

### BARseq3 Gene Module library synthesis

For fresh-frozen tissue, we rehydrated samples in 1× PBSTwG [0.1% (vol/vol) Tween-20, 100 mM glycine, 0.2 U/µL SUPERase•In RNase Inhibitor, 0.1 mg/mL yeast tRNA (Invitrogen, AM7119) in 1× PBS] for 15 min at room temperature. For fixed Robo3-CreERT2 × Ai6 samples, we washed the samples five times with PBSTwB [0.5% (vol/vol) Tween-20 (Millipore Sigma, P9416) in 1× PBS] and rehydrated them in 1× PBSTxG [0.5% (vol/vol) Triton X-100, 100 mM glycine, 0.2 U/µL SUPERase•In RNase Inhibitor in 1× PBS] for 15 min at room temperature.

We denatured SNAIL probe libraries at 90°C for 3 min immediately before hybridization. We prepared hybridization buffer with 1.5 nM per SNAIL oligo for fresh-frozen tissue (1761 gene panel) or 20 nM for fixed tissue (31 gene panel), 2× SSC (Sigma-Aldrich, S6639), 10% (vol/vol) formamide (Calbiochem OmniPur, 75-12-7), 10% (vol/vol) Triton X-100 (Sigma-Aldrich, 93443), 20 mM ribonucleoside vanadyl complex (New England Biolabs, S1402S), 0.1 mg/mL yeast tRNA (Invitrogen, AM7119), and 0.2 U/µL SUPERase•In RNase Inhibitor (Invitrogen, AM2696). We incubated the samples in hybridization buffer, covered the wells with Parafilm, placed the plates in a hybridization chamber, and incubated them at 40°C for 40 hours with gentle agitation.

After hybridization, we washed the samples twice in PBSTwR [0.1% (vol/vol) Tween-20 in 1× PBS with 0.2 U/µL SUPERase•In RNase Inhibitor] for 20 min at room temperature, and then treated them with 4× SSC in PBSTwR (4× SSC, 0.2 U/µL SUPERase•In RNase Inhibitor in PBSTwG) for 20 min at 37°C.

We prepared ligation solution [1× T4 DNA ligase buffer, 0.1 mg/mL recombinant albumin (New England Biolabs, B9200S), 0.2 U/µL SUPERase•In RNase Inhibitor, 2 U/µL T4 DNA ligase (Thermo Fisher Scientific, EL0011)] and incubated the samples for 2 hours for fresh-frozen tissue or 4 hours for fixed tissue at room temperature. We then washed the samples twice with PBSTwR for 20 min each.

We performed rolling circle amplification (RCA) in RCA solution [1× Phi29 buffer (MCLab, PP-200), 0.2 mg/mL BSA (New England Biolabs, B9200S), 0.25 mM dNTPs (Thermo Fisher Scientific, R0192), 125 µM aminoallyl-dUTP (Invitrogen, AM8439), 1 U/µL Phi29 polymerase (MCLab, PP-200), 0.2 U/µL SUPERase•In RNase Inhibitor]. We incubated the samples for 30 min at 4°C, followed by 2 hours at 30°C for fresh-frozen tissue or overnight at 30°C for fixed tissue.

We crosslinked rolonies in crosslinking solution [50 nM BS(PEG)_9_ (Broadpharm, BP-21504; stock prepared in DMSO (Thermo Fisher Scientific, D12345)), 20 mM ribonucleoside vanadyl complex in PBSTwB] for 1 hour at room temperature. We washed the samples once with 1 M Tris-HCl (pH 8.0), incubated them in fresh 1 M Tris-HCl (pH 8.0) for 30 min at room temperature, and washed them twice with PBSTwB.

### BARseq3 Sequencing

#### Sequencing primer hybridization

We prepared stripping buffer [40% (vol/vol) formamide (Calbiochem OmniPur, 75-12-7), 1% (vol/vol) Triton X-100 (Sigma-Aldrich, 93443)]. We incubated the samples in stripping buffer on a 60°C heated metal block for 5 min, allowed them to cool to room temperature, and repeated the incubation once more. We washed the samples twice with PBSTwS [2% (vol/vol) Tween-20 in 1× PBS].

We prepared sequencing primer hybridization solution [10% (vol/vol) formamide, 2× SSC] and washed the samples once in this solution. We then incubated the libraries in sequencing primer hybridization solution containing 1 µM sequencing primer for 10 min at room temperature. After hybridization, we washed the samples once in sequencing primer hybridization solution and then twice in PBSTwS.

#### Sequencing cycles

All Illumina sequencing reagents [Incorporation buffer, Incorporation Reaction Mixture (IRM), Cleavage Reaction Mix (CRM)] came from the MiSeq Reagent Nano 500 cycle Kit v2 (Illumina, MS-103-1003). For all sequencing washes performed at 60°C on a heated metal block, we cooled the samples to room temperature before exchanging solutions.

*Cycle 1:* We incubated samples twice in incorporation buffer at 60°C for 3 min each, washed them once in PBSTwS at room temperature, incubated them in iodoacetamide blocker solution [2.6 mg/mL iodoacetamide (Thermo Fisher Scientific, A39271) in PBSTwS] at 60°C for 3 min, and rinsed them in PBSTwS. We then washed the samples twice in incorporation buffer, incubated them twice in IRM at 60°C for 3 min each, and washed them four times with PBSTwS at 60°C for 3 min each. We incubated the samples with 1:100 NeuroTrace Blue Fluorescent Nissl Stain (Invitrogen, N21479) in PBSTwB for 20 min and washed them three times with PBSTwB before imaging. For Robo3-CreERT2 × Ai6 fixed tissue, we imaged endogenous ZsGreen1 fluorescence during the first sequencing cycle.

*Cycles 2–n:* We imaged ten sequencing cycles for fresh-frozen samples and five cycles for fixed tissue. We washed the samples twice in incorporation buffer, incubated them twice in CRM at 60°C for 3 min each, washed them twice with incorporation buffer, rinsed them in PBSTwS at room temperature, incubated them in iodoacetamide blocker solution [2.6 mg/mL iodoacetamide in PBSTwS] at 60°C for 3 min, and rinsed them in PBSTwS. We then washed the samples twice in incorporation buffer at room temperature, incubated them twice in IRM at 60°C for 3 min each, and finally washed them four times in PBSTwS at 60°C for 3 min each before imaging.

For experiments using the expanded 1,761-gene panel, we replaced 2.6 mg/mL iodoacetamide with 0.4% MMTS (Thermo Fisher Scientific, 23011) diluted in PBSTx (0.1% Triton X-100 in 1× PBS), and we used PBSTx instead of PBSTwS for all sequencing cycles. After the first sequencing cycle, we incubated the samples in 1:1000 DAPI (Thermo Fisher, 62248) in PBSTx for 5 min and then incubated them with 1:100 NeuroTrace Blue Fluorescent Nissl Stain in PBSTx for 20 min. We performed ten sequencing cycles for these experiments. We processed images and decoded amplicons as described previously^17^.

#### Microscopy and image acquisition

We imaged samples using a Crest X-Light V3 spinning-disk confocal mounted on a Nikon Ti2-E microscope with either a 25× silicone oil objective (NA 1.05) for Robo3-CreERT2 × Ai6 experiments or a 40× water-immersion objective (NA 1.25) for all other experiments. We excited Illumina fluorophores at 514 nm (G/T) and 640 nm (A/C). We acquired two channels simultaneously using two Hamamatsu ORCA-Fusion BT cameras synchronized by hardware TTL triggering from the microscope controller with NIS-Elements software (version 5.41.00). We separated fluorescence using a ZT405/514/635rpc dichroic with a ZET532/640m emission filter and a camera cube containing a T637/105dcbp-UT2 camera dichroic and an ET550/40+752/95m cleanup filter. We acquired z-stacks with 7 planes over 10 µm and 2× averaging. We selected imaging fields to capture the full extent of the cerebellar anlage and/or cerebellar nuclei.

### Lineage tracing with Robo3-CreERT2 × Ai6 mice

#### Tamoxifen administration

We crossed Robo3-CreERT2 mice with Ai6-ZsGreen1 reporter mice (strains listed in “Animals”). We dissolved 4-hydroxytamoxifen (Sigma-Aldrich, H6278) in corn oil at 20 mg/mL and injected it intraperitoneally into pregnant mothers on three consecutive days (E10.5, E11.5, and E12.5), using 100 mg/kg based on the mother’s weight each day. We collected embryos at E18.5 for BARseq3 analysis with the 31-gene targeted panel containing cerebellar cell-type markers.

#### Nissl stain and mounting of non-BARseq3 embryos

We fixed and embedded embryos as described in the section “Fixed tissue preparation for Robo3-CreERT2×Ai6 embryos.” We cryosectioned embryos at 30 µm onto Superfrost Plus slides and air-dried the sections at room temperature for 15 min. We then incubated the sections with NeuroTrace 435/455 Nissl stain (1:200; Thermo Fisher Scientific, N21479) for 20 min and washed them three times in 1× PBS for 5 min each. We mounted the sections with Epredia Mounting Medium (Fisher Scientific, 22-110-610) and sealed the coverslips with clear nail polish.

### Single-nucleus RNA sequencing

#### Tissue preparation

We flash-froze fresh mouse or chicken embryos in OCT and stored them at –80°C until use. We cryosectioned embryos to 100–250 µm using a cryostat onto Superfrost slides and dissected cerebellar nuclei under a dissection scope while the slide sat on a chilled metal block in a freezing CaCl_2_ bath^87^. We collected tissues of interest in 1.5 mL tubes, immediately froze them on dry ice, and stored them at –80°C until further use.

#### Nuclei isolation

We removed tissue from –80°C and immediately lysed it in 500 µL cold 0.01× lysis buffer (10 mM Tris-HCl, pH 7.4; 10 mM NaCl; 3 mM MgCl_2_; 0.1% Tween-20; 0.1% NP-40; 0.01% digitonin; 1% BSA; 1 mM DTT; 1 U/µL SUPER-ase-In RNase inhibitor). We gently triturated the tissue 10–20 times with a 1 mL pipette tip until it was mostly homogenized, incubated it on ice for 30 s, and then added 500 µL cold wash buffer (10 mM Tris-HCl, pH 7.4; 10 mM NaCl; 3 mM MgCl_2_; 1% BSA; 0.1% Tween-20; 1 U/µL SUPER-ase-In RNase inhibitor; 1 mM DTT). We mixed the sample gently and passed it through a 70 µm cell strainer into a clean microcentrifuge tube, then passed it through a 20 µm cell strainer into a second clean tube. We centrifuged the sample at 500 rcf for 5 min at 4°C, removed the supernatant, and resuspended the nuclei pellet in 1 mL chilled wash buffer (10 mM Tris-HCl, pH 7.4; 10 mM NaCl; 3 mM MgCl_2_; 1% BSA; 0.1% Tween-20; 1 U/µL SUPER-ase-In RNase inhibitor; 1 mM DTT). We repeated the 500 rcf centrifugation and wash once more, then removed the supernatant and resuspended the final nuclei pellet in resuspension buffer (1× PBS; 1% BSA; 1 U/µL SUPER-ase-In RNase inhibitor; 1 mM DTT) for downstream analysis. This protocol was adapted from ^88^.

For nuclei counting, we mixed 10 µL of the nuclei suspension with 10 µL of 0.4% trypan blue, loaded 10 µL onto a hemocytometer, and counted nuclei using standard methods. We assessed nuclei quality under high magnification to confirm intact morphology and the absence of clumps or large debris. We diluted the nuclei suspension to approximately 2000 nuclei/ µL per 10× Chromium Next GEM Single Cell 3’ Reagent Kit v4 reaction. We prepared snRNA-seq libraries according to the manufacturer’s protocol, using 11 PCR cycles for cDNA amplification and 11 PCR cycles for final amplification of gene expression libraries. We sequenced libraries on an Illumina NovaSeqX platform.

### Computational analysis

We performed all computational analyses using Python (version 3.12) and R (version 4.5).

#### Single-cell RNA sequencing data analysis Data processing

We processed demultiplexed reads with 10X Genomics CellRanger count (CellRanger v7.2.0) using default parameters. We aligned RNA-seq reads to the mm10 reference genome for mouse and GRCg7b (galGal7) for chicken datasets. We processed count data with the Seurat R package (v5.0.1). We filtered cells to include those with counts above 2,000 and below 20,000 for the chicken dataset or above 2,000 and below 10,000 for the mouse dataset, and we removed all cells with ribosomal or mitochondrial transcripts.

#### Single-cell dataset integration and cell type classification

<u>All upper rhombic lip (**Figs. 1, 2**):</u> We subsetted the Butts et al. dataset^40^ to clusters that were coarsely *En1*/*En2*+ and negative for Hox gene expression, according to their description of cerebellar lineages. We selected non-cycling neurons by excluding *Mki67*+ and *Top2a*+ cells. We used FastMNN^89^ with the top 5,000 variable genes from the SeuratWrappers package^90^ to integrate across the “Samples” variable, which encoded separate dissections of developmental tissues.

We clustered the mnn embedding with Seurat Louvain clustering using 20 nearest neighbors and resolution 2. We manually merged initial clusters when they met the following criteria: (1) Pearson correlation between clusters > 0.8 for top marker genes; (2) similar marker-gene expression patterns; and (3) representation of the same developing cell type across adjacent timepoints, with observed differences attributed to developmental progression rather than distinct cellular identity.

<u>All excitatory cerebellar nuclei (**Figs. 2, 3**):</u> From the cell types found in the RL-derived excitatory neurons object described above, we subsetted the clusters assigned to the cerebellar nuclei based on absence of *Zfhx3* and presence of *Meis2* expression. We then combined this dataset with a manually curated E17.5 snRNA-seq dataset and a published adult excitatory cerebellar nuclei dataset ^3^ by first subsetting to genes shared between datasets. We also removed sex-specific genes, including "Xist", "Tsix", "Kdm6a", "Eif2s3x", "Kdm5c", "Ddx3x", "Usp9x", "Bcor", "Car5b", "Smcx", "Shroom4", "Mecp2", "Ddx3y", "Eif2s3y", "Kdm5d", "Uty", "Usp9y", "Jarid1d", "Rbmy", "Zfy1", "Zfy2", "Sry", "Tspy", "Pramel1", "Tmsb4y". We performed SCTransform^91^ on the combined object and ran FastMNN with 2,000 features, integrating over separate tissue dissections, to generate excitatory embeddings spanning development to adulthood.

We clustered the mnn embedding with Seurat Louvain clustering using 20 nearest neighbors and resolution 1.2. We manually merged clusters when they met the criteria described above. We assigned clusters to nuclei based on their transcriptomic similarity to adult cerebellar nuclei and their spatial proximity to the adult nuclei in embedding space. We also used known marker gene expression to annotate clusters (for example, *Lmx1a* for the medial nucleus and *Lhx9* for interposed and lateral nuclei).

<u>All ventricular zone (**Figs. 1, 5**):</u> We subsetted five published datasets^3,27–30^ to clusters labeled as inhibitory neurons in the developing cerebellum. We subsetted our manually curated E17.5 dataset to Gad1+ clusters after subsetting the objects to shared genes. We used FastMNN with the top 500 variable features to integrate samples across datasets.

We clustered the mnn embedding with Seurat Louvain clustering using 20 nearest neighbors and resolution 2. We manually merged initial clusters when they met the criteria described above.

<u>Excitatory progenitor cell types (RL, **Fig. 4**):</u> We subsetted the Butts et al. dataset^40^ to clusters that were *En1*/*En2*+ and *Hox*– and had *Mki67* > 1, which labeled cycling progenitors, as described in their paper. To find unbiased clusters from this dataset, we integrated the dataset using FastMNN with 2,000 features across “Samples” and clustered in this mnn space.

#### Pseudotime analysis (Supplementary Fig. 5)

We inferred developmental trajectories for excitatory postmitotic cerebellar nuclei neurons using a Seurat-to-Monocle3 workflow. We first subsetted the combined multi-timepoint dataset to the clusters of interest, and excluded adult cells in the analysis. We then converted the subset to a Monocle3 cell_data_set, transferred the Seurat UMAP coordinates, and stored cluster labels in colData. We learned the trajectory graph with learn_graph(), and ordered cells in pseudotime using a designated root cluster as the rhombic lip (RL).

For each trajectory, we plotted pseudotime together with smoothed expression profiles for selected genes of interest, including *Lhx9*/*Pax6*, *Olig2*/*Irx3*/*Lmx1a*, and *Prox1*/*Zfpm2*. Gene expression trends were smoothed with LOESS (locally estimated scatterplot smoothing), which is in the Monocle3 package.

#### Excitatory progenitor classifier implementation (Fig. 4)

We trained the classifier on postmitotic RL-derived excitatory cells to capture mature cell-type signatures, and then applied it to RL progenitors to predict underlying developmental fate: We trained a multinomial lasso regression classifier in the R package glmnet v5.0 to predict cycling progenitor (RL) wave identity^92^. To do this, we first randomly split postmitotic cells from the postmitotic RL-derived excitatory cell dataset from E12.5–E13.5 into training (50%) and test (50%) sets. We subsetted all data to transcription factors. We identified differentially expressed transcription factors as shared differentially expressed genes between RL waves and variable genes of RL progenitor cells, and we used these features to train the classifier.

We normalized and scaled RL expression values in Seurat, and we fit a multinomial logistic regression model (alpha = 0 or 1) using the selected features. We evaluated model performance on the held-out test set.

We found a different confidence threshold for each cell type by finding the highest threshold at which the 90% of the test data was accurately classified (**Extended Data Fig. 6C,D**). We then applied the trained classifier to the RL subset by normalizing and scaling the query dataset, extracting scaled expression values for the shared feature set, and constructing a gene-by-cell matrix spanning the training features. We applied confidence thresholds to cycling progenitors, and we retained these cells for downstream analysis and/or colored them in relevant plots, while we excluded or colored lower-confidence cells in gray.

To evaluate the concreteness of our classifications, we performed permutation test for both excitatory lineage branches (early vs late) and all early rhombic lineage neurons (Med^early^ vs IntA/Lat vs IntP vs extracerebellar-fated neurons). First, we permuted labels of each cell type in the training data and saw poor probability distributions and low confidence of progenitor classification (**Extended Data Fig.6E**). Next, we permuted each gene name in the progenitor data, which lead to very low classification efficiency. (**Extended Data Fig. 6F**)

#### Chicken vs. mouse comparison (Fig. 4)

We extracted SLC17A6+ cells that clustered separately from other populations from the chicken snRNA-seq dataset. Among these, we selected clusters that were MEIS2+ and ZFHX3– for comparative analysis with mouse cerebellar nuclei. We assessed cross-species similarity using MetaNeighbor, based on shared highly variable genes restricted to one-to-one orthologs between chicken and mouse.

#### RA-FGF8 Module Scores (Extended Data Fig. 8A-D)

We quantified RA-positive and FGF8-positive transcriptional programs using Seurat module scoring. We defined two gene sets: an RA-positive program (*Nr2f2, Nr2f1, Meis2, Meis1, Crabp1*) and an FGF8-positive program (*Cnpy1, En1, En2*). Prior to scoring, each gene set was restricted to genes present in the RNA assay, and any missing genes were recorded. Module scores were calculated with Seurat’s AddModuleScore() using 20 matched control genes per feature set and a fixed random seed for reproducibility. This analysis was performed on mouse samples from E12.5, E13.5, and E14.5, as well as on chicken E6/7 samples using orthologous genes.

### BARseq3 data analysis

We analyzed all BARseq3 data using Scanpy (version 1.8)^93^ in Python (version 3.12).

#### Integrated cross-timepoint BARseq3 data (Figs. 1 & 5)

We selected representative slices from each timepoint that contained all relevant cell types and areas of interest, and we concatenated AnnData objects for downstream analysis. We computed quality-control metrics using Scanpy and then filtered out cells with fewer than 5 detected counts and over 400 counts and genes detected in fewer than 10 cells. We normalized expression values per cell using total-count normalization, log-transformed them, and scaled them with a maximum value of 10. We performed principal component analysis using the ARPACK solver and then applied Harmony integration (version 1.2)^94^ across library_id to correct for batch effects. We constructed a 6-nearest-neighbor graph using the Harmony-corrected PCA embedding and used UMAP for visualization. We removed clusters containing fewer than 10 cells. We performed unsupervised clustering with Leiden clustering and manually annotated cluster identities based on canonical marker-gene expression. We assigned Leiden clusters to broad cell-type categories, including cerebellar nuclei, granule cells, molecular layer interneurons, glia/oligodendrocytes, midbrain-fated cells, choroid plexus, and outside cerebellar populations.

To transfer these annotations to additional slices, we used a manually labeled reference slice as the training set and unlabeled slices from the same timepoint as query data. We restricted reference and query datasets to shared genes, normalized them by total counts, and log-transformed them. We identified highly variable genes from the reference dataset and used them to define a shared feature space. We scaled query data using the mean and standard deviation calculated from the reference data and performed principal component analysis on the reference dataset. We projected query cells into the same PCA space using the reference PCA loadings. We transferred cell type labels using a k-nearest-neighbor classifier in PC space (k = 15, distance-weighted voting), and we assigned each query cell the label with the highest predicted probability. We visualized predicted labels in UMAP and spatial coordinates using a consistent color scheme and category order across slices.

We used excitatory and inhibitory cerebellar nuclei cell types defined here in **Fig. 5**. **Excitatory cerebellar nuclei cell type classification (Fig. 3)**

We normalized spatial transcriptomic data in Scanpy and stored raw counts before preprocessing. We normalized expression values per cell, log-transformed them, regressed for total counts, and scaled them with a maximum value of 10. We performed principal component analysis on the transcriptomic data and then applied Harmony integration across library_id to account for library-specific variation. To define spatially informed clusters, we constructed an expression-based nearest-neighbor graph from the Harmony-corrected PCA embedding and combined it with a spatial neighbor graph computed using Squidpy (version 1.8)^95^. We integrated the two graphs as a weighted sum, with transcriptomic similarity weighted more heavily than spatial proximity (typically 80%-90% transcriptomic weightage, see code), and then performed Leiden clustering on the resulting hybrid graph to identify major spatial cell populations.

We separated excitatory cerebellar nuclei from other populations based on their spatial position within the cerebellar anlage and expression of canonical markers such as *Slc17a6* and *Meis2*. We then subsetted a broad cerebellar cluster and reclustered it using the same hybrid expression–spatial framework to refine excitatory neuronal subtypes. We removed clusters containing fewer than 10 cells. We merged and defined these clusters based on the best fit across timepoints, as specified below.

#### Cross-timepoint correspondence analysis (Fig. 3)

We assessed cell type correspondences across developmental stages using MetaNeighborUS (pymn, version 1.1)^96^. For each pairwise comparison, we concatenated AnnData objects from two timepoints and retained cell-type labels and study identifiers in the metadata. We computed correspondence scores using highly variable genes identified within the combined dataset, depending on the analysis. We ran MetaNeighborUS in both one- vs-best and full pairwise modes to generate AUROC-based similarity scores between cell types across timepoints. We then extracted scores for the two timepoints of interest and visualized them as heatmaps of cross-timepoint correspondence. Clusters merged and matched using a greedy algorithm, in which we first identified the pair of clusters with the highest MetaNeighborUS correspondence score and assigned them as a match. We then removed that pair from further consideration and repeated the process with the remaining clusters, always selecting the next highest-scoring pair. This approach continued until all clusters were assigned or no additional pair met the matching criteria.

#### Spatial cell type projection (Fig. 3)

To quantify the spatial distribution of excitatory neuronal subtypes along the mediolateral axis, we projected cells onto a line segment defined by two spatial coordinates and parallel to the mediolateral axis of the excitatory cerebellar nuclei. For E11.5 to E13.5, we manually selected points at the edge of the rhombic lip and the edge of the cerebellar nuclear transitory zone. For E14.5 to E17.5, we selected points across all mediolaterally arranged cerebellar nuclei. We assigned each cell of interest a scalar position along the segment by orthogonal projection and clipped projections to the segment boundaries to restrict analysis to the region of interest.

We discretized projected coordinates into 60 equally spaced bins along the axis. For each bin, we computed the fraction of cells belonging to each excitatory group from the cell types defined above. We assigned zero values to bins containing no cells. To reduce local variation, we smoothed the resulting fraction profiles using a centered rolling mean with a window size of 10 bins. We normalized curves by the sum of values across the mediolateral axis.

For inhibitory neurons (**Fig. 5**), we applied the same procedure to lines spanning the dorsoventral axis, restricting the analysis to cells within 250 µm on either side of each line and grouping cells by broad cerebellar nuclei cell types.

#### scRNAseq and BARseq3 correspondence (Fig. 3)

We quantified transcriptional correspondence between scRNAseq reference populations and BARseq3 cell populations using a pseudobulk correlation–AUROC procedure. At each developmental stage, we restricted both datasets to shared genes and excluded cell populations represented by fewer than 10 cells. We calculated the mean expression profile of each annotated cell population in each dataset and selected up to 500 genes with the greatest variation across population-level expression profiles. For each BARseq3 cell, we calculated the Spearman correlation between its expression profile and the pseudobulk expression profile of each scRNAseq reference population. For every reference–BARseq3 population pair, we then calculated a one-versus-rest area under the receiver operating characteristic curve (AUROC), using the correlation score to distinguish cells belonging to the indicated BARseq3 population from all other BARseq3 cells. An AUROC of 0.5 indicates no discriminatory correspondence, whereas values approaching 1 indicate stronger transcriptional correspondence between the reference and BARseq3 populations. E15.5 and E17.5 BARseq3 populations were compared with the E16.5 scRNAseq reference because a stage-matched reference dataset was unavailable.

#### Excitatory and inhibitory integration analysis (Fig. 5)

We identified excitatory neurons from spatial clustering at each developmental stage. We identified i1 inhibitory neurons as clusters expressing *Sox14*, *Dmbx1*, and *Zfhx3*. We identified i2/3 neurons as *Pax2*+ but could not distinguish them from other cerebellar interneurons because the BARseq3 panel lacked specific marker genes.

To evaluate integration, we quantified the local cellular environment surrounding each excitatory cerebellar nuclei neuron by identifying its 20 nearest neighboring nuclei cells and calculating the ratio of inhibitory (independently for i1 and interneurons) to excitatory neighbors. We performed this analysis at each developmental stage and compared the resulting neighborhood composition to the adult pattern to determine when local spatial organization became adult-like. We restricted the analysis to neighbors within 120 µm of each excitatory neuron.

#### Adult cerebellar nuclei cell type distribution (Fig. 6)

We defined adult cerebellar nuclei subdivisions (Lat, IntA, IntP, MedL, MedC, MedDL) based on Nissl images, cytoarchitectonic boundaries based on cell density, and annotations in the Paxinos and Franklin atlas^97^. We defined coarse cell types based on excitatory cerebellar nuclei markers (*Slc17a6*, *Meis2*) and inhibitory cerebellar nuclei markers (i1: *Sox14*; i2/3: *Pax2*). The proportion of i1 and i2/3 neurons within these subnuclei was computed.

### snATACseq analysis (Extended Data Fig. 7)

We processed snATAC-seq data from the E12.5, E13.5, and E14.5 datasets of ^28^ using a pipeline adapted from their code. Fragment files were converted to Arrow files with the mm10 genome, requiring a minimum TSS enrichment of 5 and at least 1,250 fragments per cell. Doublets were inferred but retained. We created an ArchR project and performed iterative LSI on the TileMatrix, followed by Seurat-based clustering and UMAP visualization.

To annotate cell identities, we examined GeneScoreMatrix profiles for marker genes including *Olig2, Zfhx3, Meis2, Ntng1, Lmx1a, Slc17a6, Gad1, Robo3*, and *Atoh1*. We then subsetted clusters of interest and re-ran LSI, clustering, and UMAP to refine lineage-specific population structure, including a higher-resolution analysis of the cerebellar nuclear lineage subset. We added a peak set and peak matrix, and integrated the scRNAseq reference using ArchR’s addGeneIntegrationMatrix() to transfer RNA-defined branch annotations to ATAC cells.

We also assessed TSS enrichment and fragment counts as quality-control metrics. Finally, we annotated motifs using the cisBP database, computed motif deviation scores with addDeviationsMatrix(), and evaluated RARE motif activity across ATAC clusters. Motif marker analysis and footprinting were performed for retinoic acid receptor–related motifs in selected groups, including IntP, IntA, and Lat, to assess lineage-specific retinoic acid regulatory activity.

### Robo3-CreERT2 x Ai6 Zsgreen1 analysis (Fig. 6)

We identified candidate *ZsGreen1*+ cells as cells with a maximum fluorescence intensity greater than 1,000 and generated 100 µm bounding boxes around each candidate cell. We randomly ordered the cells and asked three expert annotators to classify each candidate as a true or false positive blinded to the location of each cell. We included cells in the final analysis when at least two of the three annotators classified them as *ZsGreen1*+. Annotator results can be found in **Extended Data Table 2**, with accompanying cell data in the Zenodo data repository.

We classified cells as excitatory or inhibitory based on marker gene expression (*Slc17a6*, *Meis2* for excitatory; *Sox14* and *Dmbx1* for i1 inhibitory; *Pax2* for i2/3 interneurons) and assigned them to cerebellar nuclei subdivisions (Lat, IntA, IntP, MedL, MedC/DL) based on marker gene expression, spatial clustering (see above), and cytoarchitectonic boundaries visible in Nissl staining. We quantified the fraction of ZsGreen1+ excitatory and inhibitory neurons for each nucleus subdivision.

## DATA AND CODE AVAILABILITY

Website for interactive data objects is available at https://mmganant.github.io/CNdev_website/. Code for all analyses is available at https://github.com/mmganant/CNdev_analysis. Processed data are available at https://zenodo.org/records/21796565. Raw snRNAseq FASTQ files are available at BioProject Accession PRJNA1531022. Raw BARseq3 imaging data is available upon request.

## ACKNOWLEDGEMENTS

We thank members of the Kebschull and Fan labs for helpful discussions, feedback on the project, and technical assistance, and Liqun Luo, Alex Kolodkin, and Reza Kalhor for feedback on the manuscript. We thank Huihui Qi for annotations of Zsgreen1 fluorescent cells in our transgenic mice experiments. We thank Kevin Wright (Oregon Health and Science University) for providing Robo3-CreERT2 mice.

## FUNDING

This work was supported by a Packard Fellowship, a Klingenstein-Simons Fellowship, a Sloan Fellowship, and a Pershing Square MIND prize to J.M.K., a National Institute of General Medical Sciences award R35GM142889 to J.F., a Kavli Distinguished PhD Fellowship to M.M.G.A., and an NSF GRFP to C.S.. M.L.P. was supported by National Institute of General Medical Sciences award R35GM155257.

## AUTHOR CONTRIBUTIONS

M.M.G.A. and J.M.K. conceived the project and designed the experiments. M.M.G.A. performed animal dissections, tissue collection, lineage-labeling, library preparation, and sequencing experiments. M.L. administered 4-hydroxytamoxifen for Robo3-CreERT2 experiments and set up timed pregnancies. M.M.G.A. and E.C.Z. imaged BARseq3 experiments. C.S. assisted with running the 10× Genomics Chromium system for snRNA-seq experiments. D.Z.F.G. assisted in generating oligonucleotide probes for BARseq3 experiments. M.L.P. assisted in the chicken experiments. M.M.G.A. performed all computational analyses and interpreted the results. M.M.G.A. prepared the figures. M.M.G.A. and J.M.K. wrote the manuscript, with assistance from J.F.. J.M.K. and J.F. funded all experiments. All authors reviewed and approved the final manuscript.

## DECLARATION OF INTERESTS

The authors declare no competing interests.

## AI STATEMENT

We used artificial intelligence-assisted tools to help edit text, improve clarity, and assist with code drafting during manuscript preparation and data analysis. The authors reviewed and verified all scientific interpretations, analyses, figures, and conclusions. We did not use any AI tool as a substitute for author judgment, experimental validation, or independent data interpretation.

## EXTENDED DATA FIGURES

**Extended Data Figure 1.**
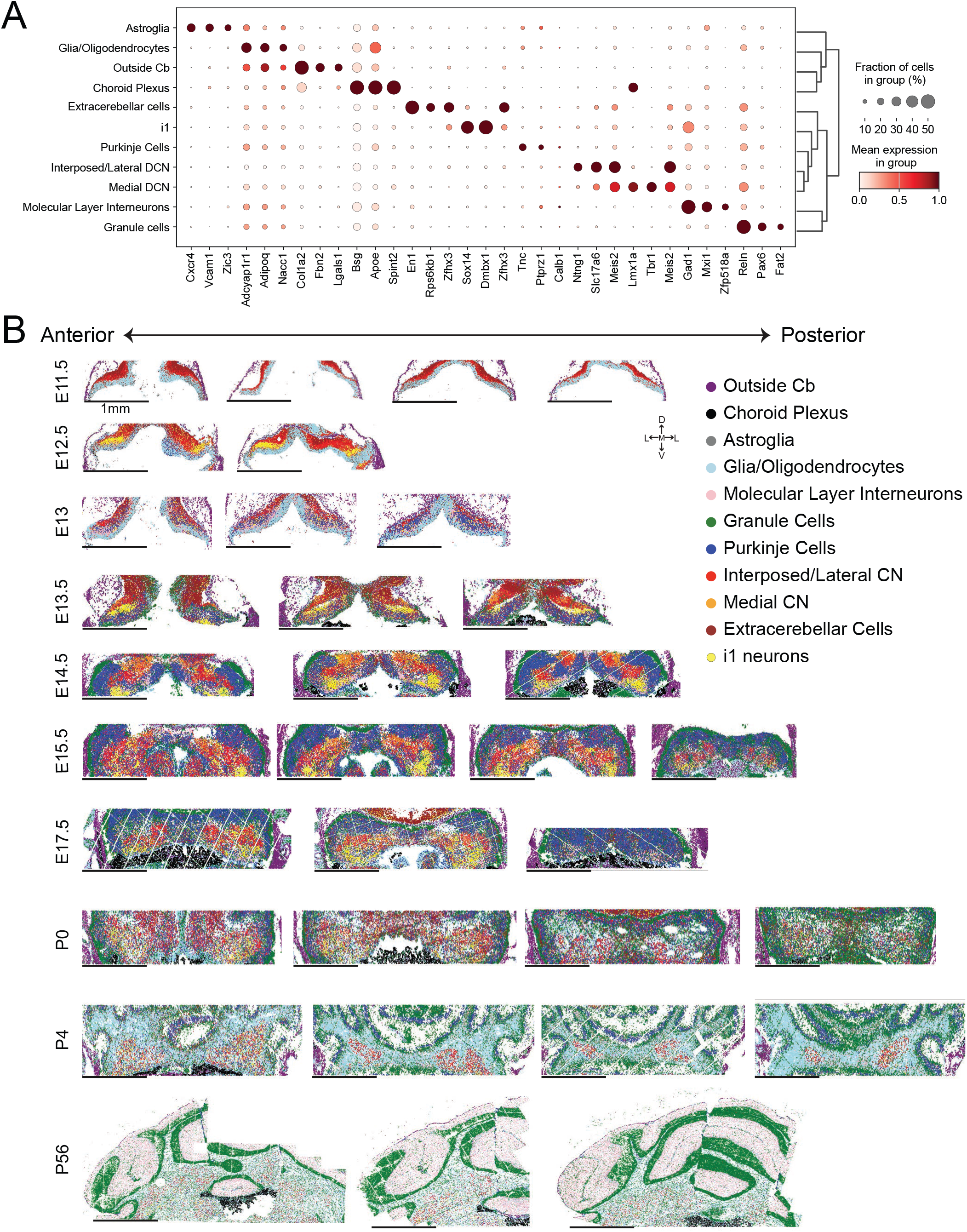
All BARseq3 sections. **(A)** Dot plot of marker genes defining cell types derived from integrated BARseq3 datasets across timepoints. **(B)** All 33 BARseq3 sections, arranged from anterior to posterior, across developmental timepoints and colored by integrated cell type.

**Extended Data Figure 2.**
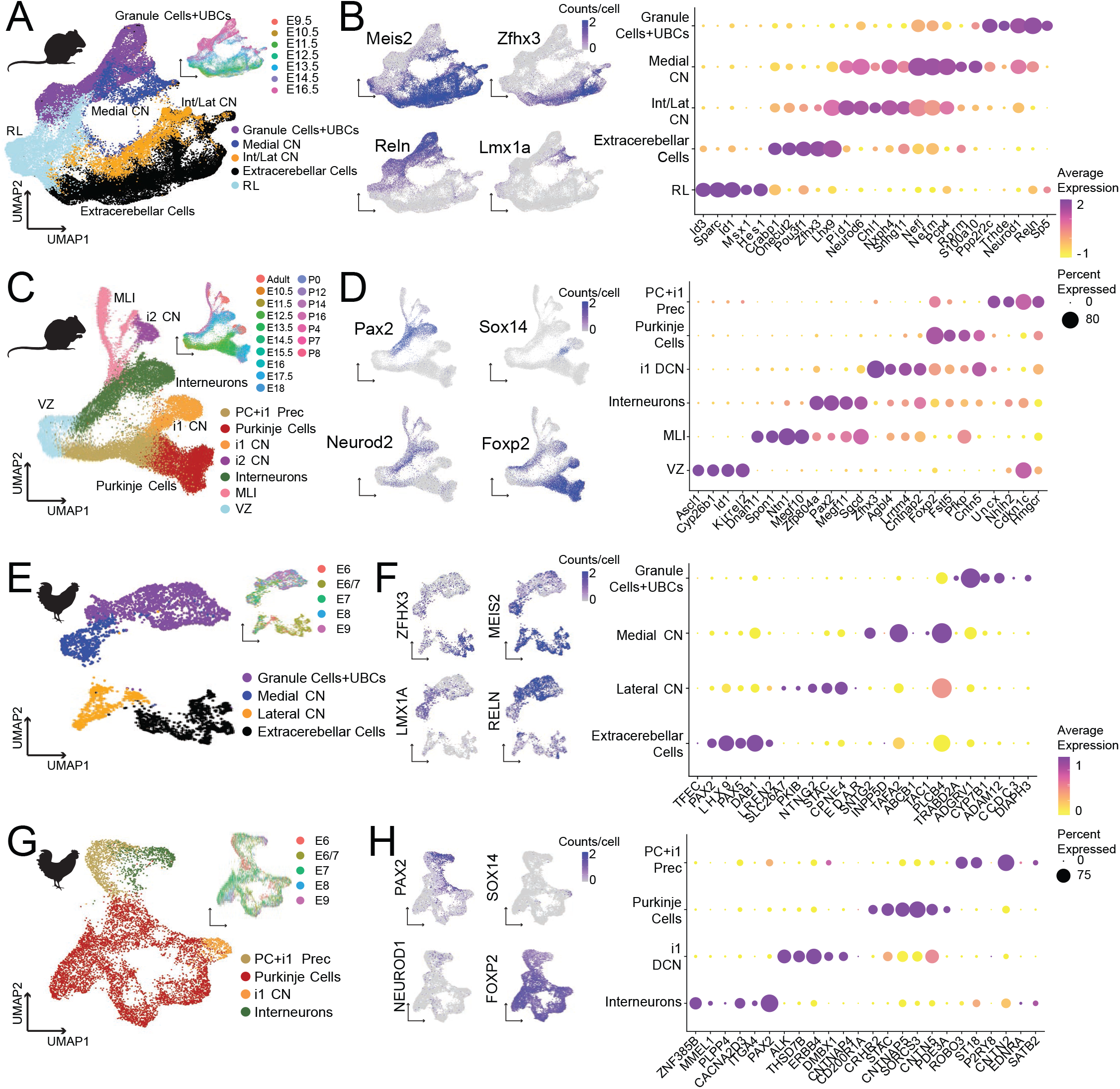
scRNAseq dataset marker gene expression. **(A)** UMAP embedding of upper rhombic lip derived postmitotic mouse excitatory neurons from embryonic day (E) 9.5 to E16.5, revealing major cell types including granule cells and unipolar brush cells (UBCs), medial and interposed/lateral (Int/Lat) cerebellar nuclei cells, extracerebellar cells, and early postmitotic precursors at the rhombic lip (RL). **(B)** Expression of canonical marker genes for each population (left) and dot plot of differentially expressed genes (right) across all excitatory neuron types. **(C)** UMAP embedding of ventricular zone derived mouse inhibitory cerebellar neurons from E10.5 to adult, revealing major cell types including Purkinje cells, cerebellar nuclei inhibitory neuron subtypes (i1, i2, i3), intermediate Purkinje cell and i1 precursors (PC+i1 Prec), cerebellar cortical interneurons, molecular layer interneurons (MLIs), and early postmitotic precursors at the ventricular zone (VZ). **(D)** Expression of canonical marker genes for selected populations (left) and dot plot of differentially expressed genes (right) across all inhibitory neuron types. **(E)** UMAP embedding of chicken excitatory neurons from E6 to E9, showing conserved cell types including granule cells, chicken medial and lateral cerebellar nuclei neurons, and extracerebellar neurons. **(F)** Expression of orthologous marker genes in chicken (left) and dot plot of differentially expressed genes (right), demonstrating similar expression patterns to mouse. **(G)** UMAP embedding of chicken inhibitory neurons from E6 to E9, identifying conserved cell types including Purkinje cells, i1 cerebellar nuclei neurons, their precursors (PC+i1 Prec), and interneurons. **(H)** Expression of orthologous marker genes in chicken (left) and dot plot of differentially expressed genes (right), demonstrating expression patterns similar to mouse.

**Extended Data Figure 3.**
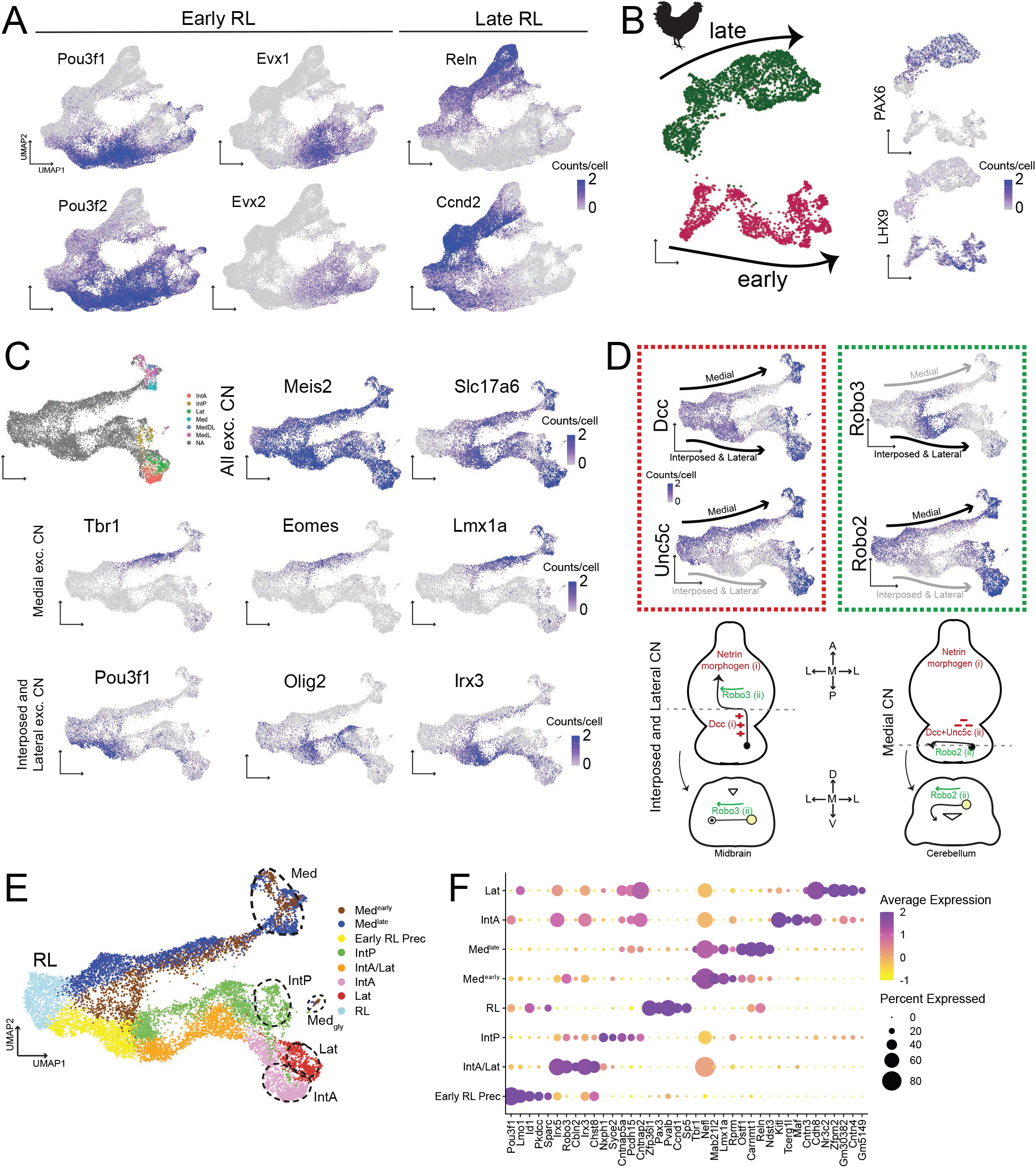
Early and late rhombic lip waves and excitatory cerebellar nuclei marker gene expression. **(A)** UMAP embedding of the mouse upper rhombic lip Atoh1-lineage scRNAseq dataset, showing marker genes for early and late waves. **(B)** UMAP embedding of the chicken excitatory scRNAseq dataset, showing early and late branches with marker genes LHX9 and PAX6. **(C)** UMAP embedding of mouse integrated excitatory cerebellar nuclei scRNAseq data, colored by adult excitatory cerebellar nuclei defined in ^3^ (top left). Expression of marker genes for the cerebellar nuclei, including *Meis2* and *Slc17a6*, as well as markers specific to the medial and interposed/lateral lineages. (**D**) Expression of axon guidance genes in the early and late cerebellar nuclei branches, which may be indicative of their projection routes out of the cerebellum: While the excitatory projections of the interposed and lateral cerebellar nucleus leave the cerebellum via the ipsilateral superior cerebellar peduncle and cross over the ventral midline in the midbrain in a Robo3 dependent manner, the projections of the medial nucleus uniquely leave the cerebellum by first crossing the midline within the cerebellum via the Hook bundle ^7,98^. High Unc5c expression and lack of Robo3 expression in the medial branch correlate with this unique path and might point to a mechanism for this rostral midline crossing. **(E)** Excitatory cerebellar nuclei scRNAseq dataset, colored by cell type, with adult nuclei defined by ^3^ indicated. **(F)** Differentially expressed genes for the cell types shown in (**E**), excluding adult cells.

**Extended Data Figure 4.**
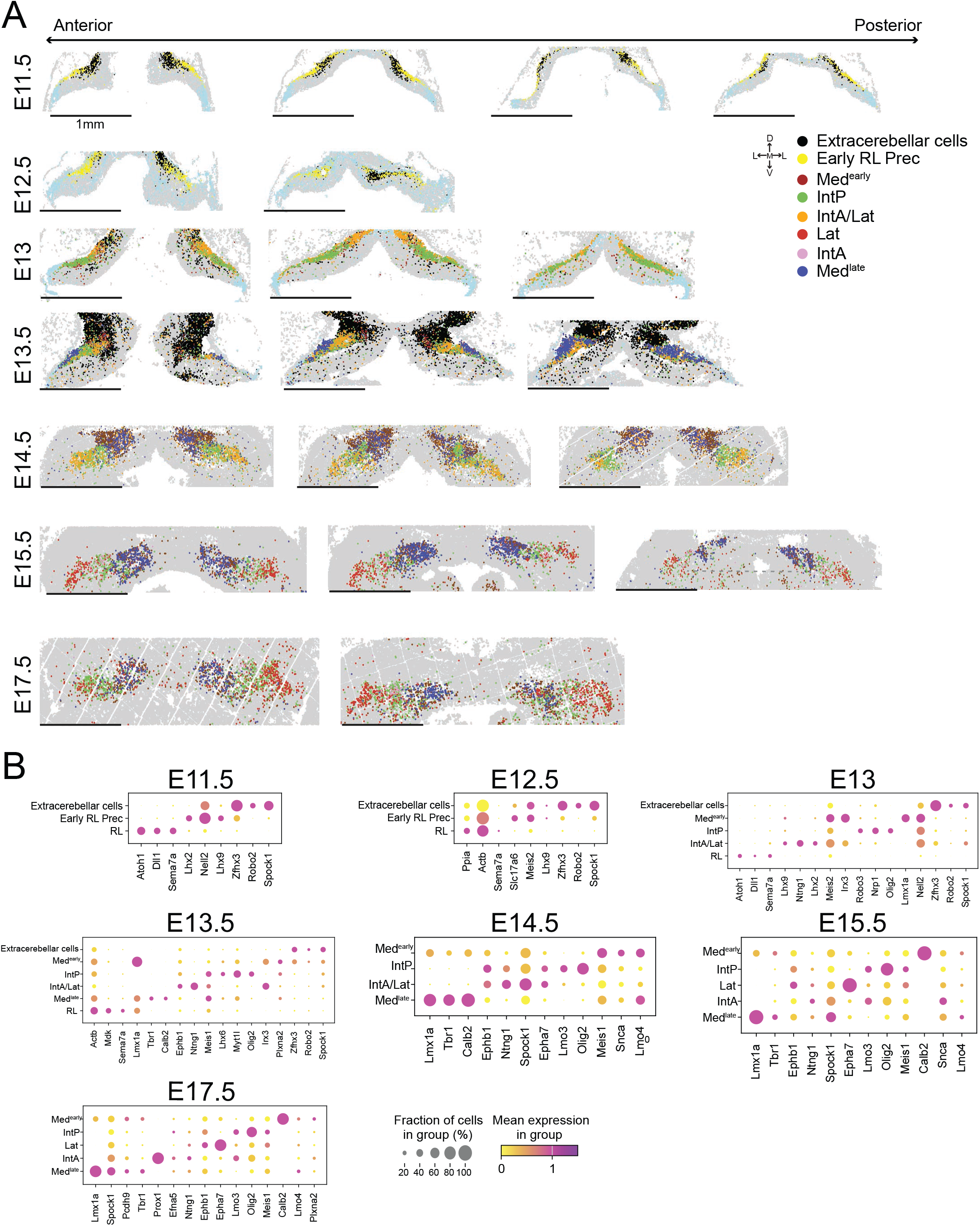
Excitatory cerebellar nuclei cell types in all BARseq3 sections. **(A)** BARseq3 sections spanning the anterior to posterior axis of the cerebellar anlage across developmental timepoints, colored by cell type. Grey indicates cells that are not excitatory cerebellar nucleus neurons, extracerebellar-fated neurons, or cells outside the cerebellum. Sections with fewer than 100 excitatory cerebellar nucleus neurons were removed from this analysis. Lat, lateral nucleus. **(B)** Dot plots of marker gene expression defining the major excitatory cerebellar nuclei cell types at each analyzed developmental timepoint.

**Extended Data Figure 5.**
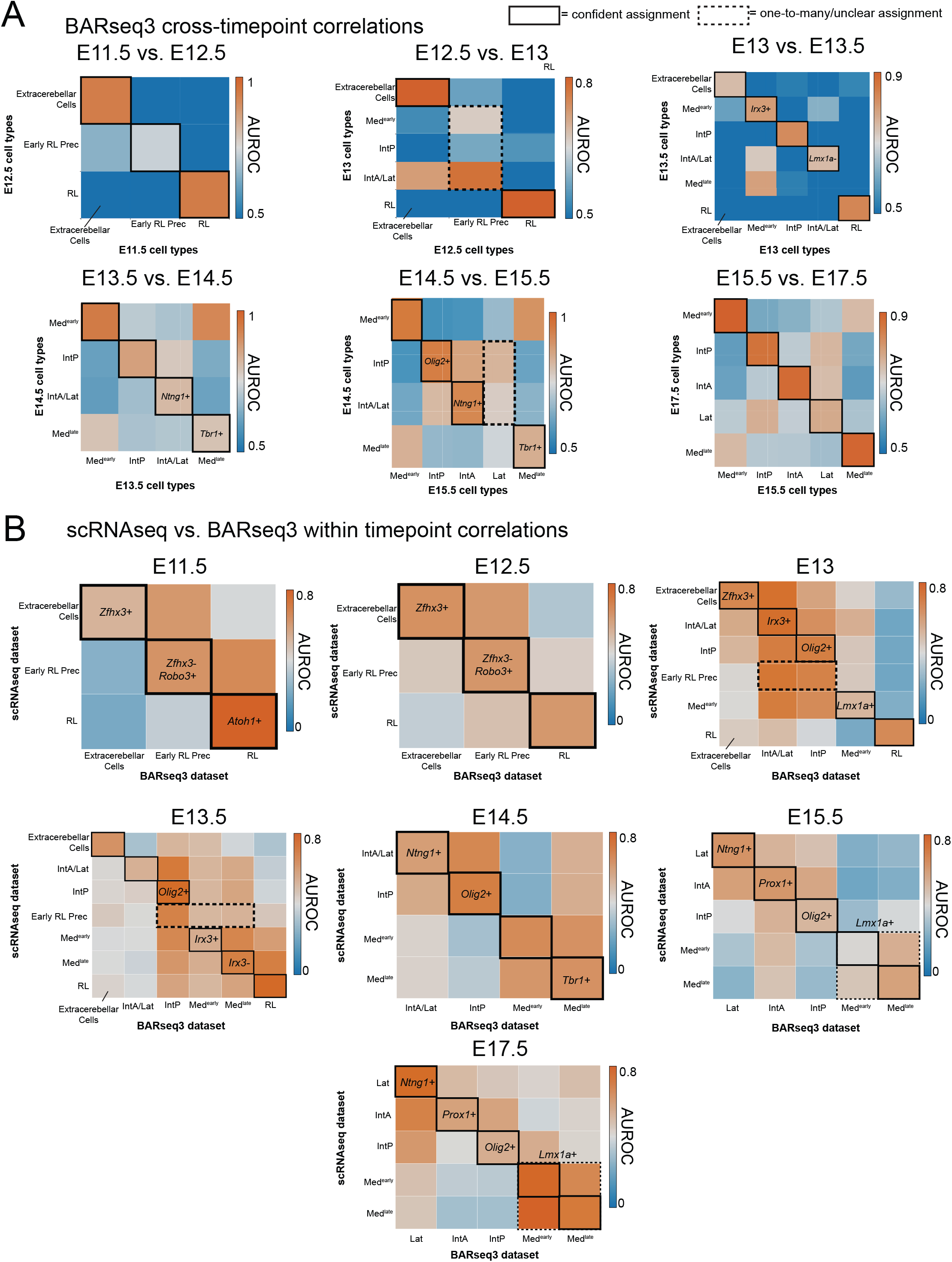
BARseq3 cross-timepoint correlations and BARseq3–scRNAseq correlations. **(A)** MetaNeighbor AUROC of BARseq3 excitatory cerebellar nuclei cell types across pairs of adjacent timepoints. **(B)** Custom AUROC (see *Methods*) of BARseq3 excitatory cerebellar nuclei cell types with matched timepoint scRNAseq excitatory cerebellar nuclei cell types.

**Extended Data Figure 6.**
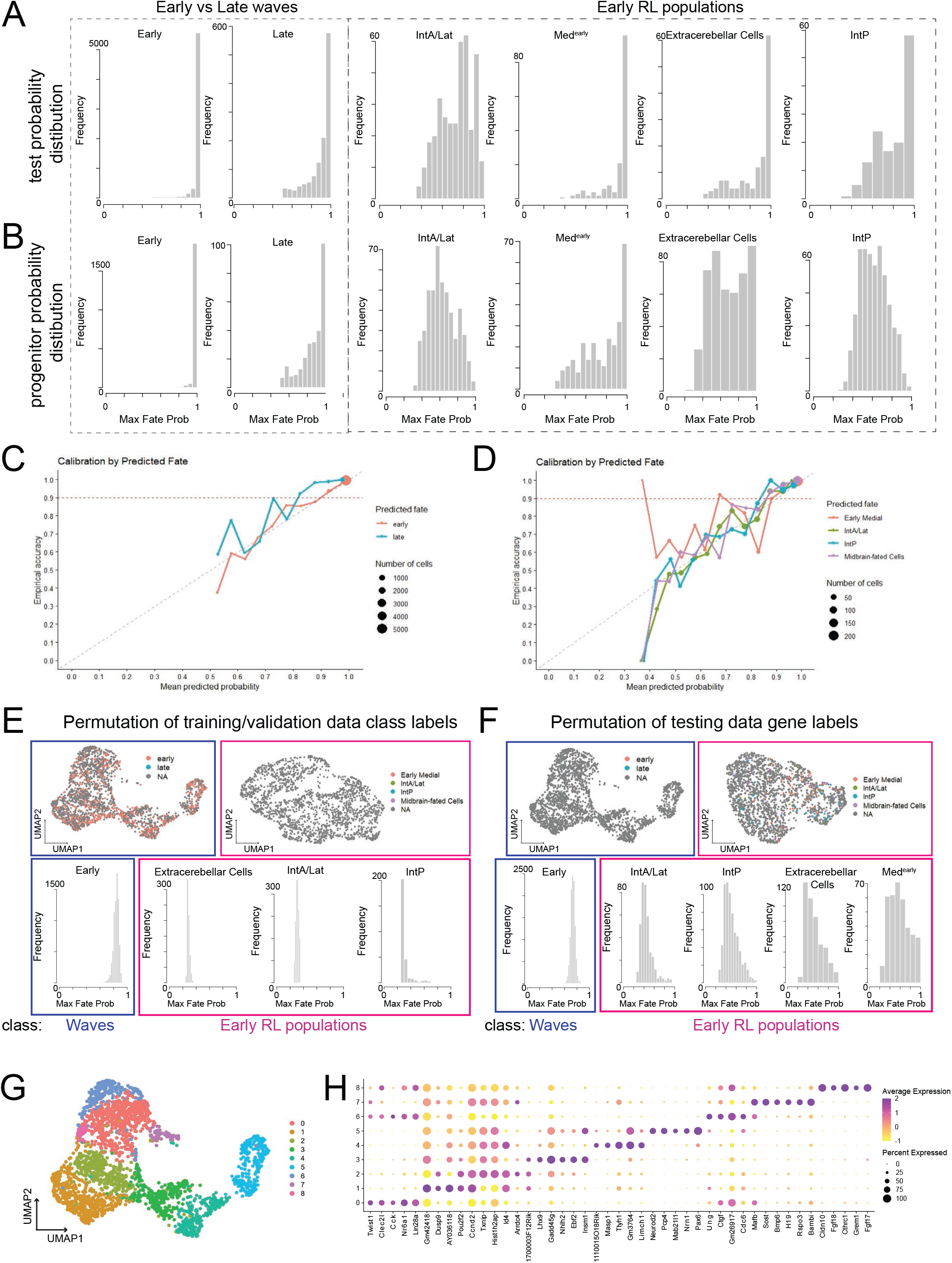
Quality control of classifiers and cycling progenitor transcriptional heterogeneity. **(A)** Histograms showing the distribution of confidence in assigning cells from a held-out postmitotic excitatory neuron test set to classifiers trained on early versus late rhombic lip (left) and early rhombic lip cerebellar nuclei cell types (right). **(B)** Histograms showing the distribution of confidence in assigning cycling progenitors to classifiers trained on early versus late rhombic lip (left) and early rhombic lip cerebellar nuclei cell types (right). **(C-D)** For held out data, line plot showing the mean probability of assignment to correct class of either **(C)** early versus late rhombic lip or **(D)** early rhombic lip cerebellar nuclei cell types. Thresholds were calculated based on 90% empirical accuracy for each cell type independently. **(E)** Classifier performance of progenitors when postmitotic excitatory neuron training/validation class labels are shuffled, with UMAP embeddings (top) and assignment probability distibutions per class (bottom), for both early versus late rhombic lip or early rhombic lip cerebellar nuclei cell types. **(F)** Same as **(E)** but shuffling gene labels of cycling progenitor test datasets. **(G)** Unsupervised clustering of excitatory rhombic lip-derived cycling progenitors. **(H)** Differentially expressed genes across clusters from **(G)**.

**Extended Data Figure 7.**
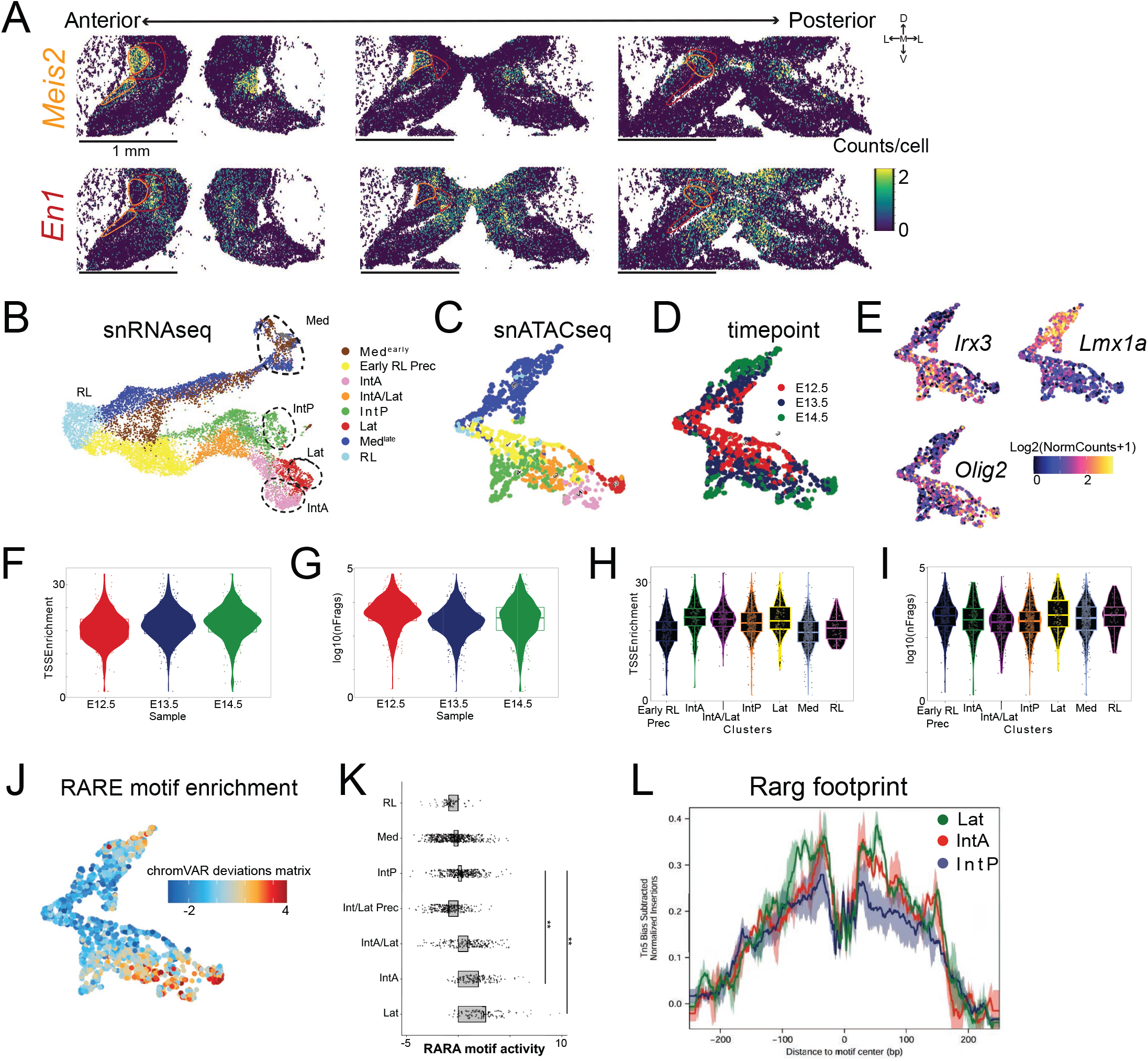
Spatial gene expression and snATACseq analysis of RA- and FGF8-associated programs that distinguish IntP from IntA/Lat. **(A)** Expression of *Meis2*, enriched in the RA pathway, and *En1*, enriched in the FGF8 pathway, showing inverse spatial expression patterns in E13.5 mouse nuclear transitory zone, as seen in BARseq3 anterior-to-posterior sections. **(B)** Excitatory cerebellar nuclei scRNAseq dataset colored by cell type. **(C)** Label transfer of scRNAseq dataset cell types onto the snATACseq embedding. **(D)** snATACseq embedding colored by timepoint. **(E)** snATACseq embedding colored by gene score for marker genes of different cerebellar nuclei populations from scRNAseq. **(F-I)** Quality control of the snATACseq dataset, showing transcription start site (TSS) enrichment across **(F)** timepoints and **(H)** cell types, and number of unique fragments across **(G)** timepoints and **(I)** cell types. **(J)** Enrichment of the RARE motif, the target of RA signaling, in the snATACseq embedding. **(K)** Quantitative evaluation of RARE motif enrichment across excitatory cerebellar nuclei cell types. **(L)** Footprint analysis of Rarg shows a larger peak-to-trough footprint for IntA and Lat compared with IntP.

**Extended Data Figure 8.**
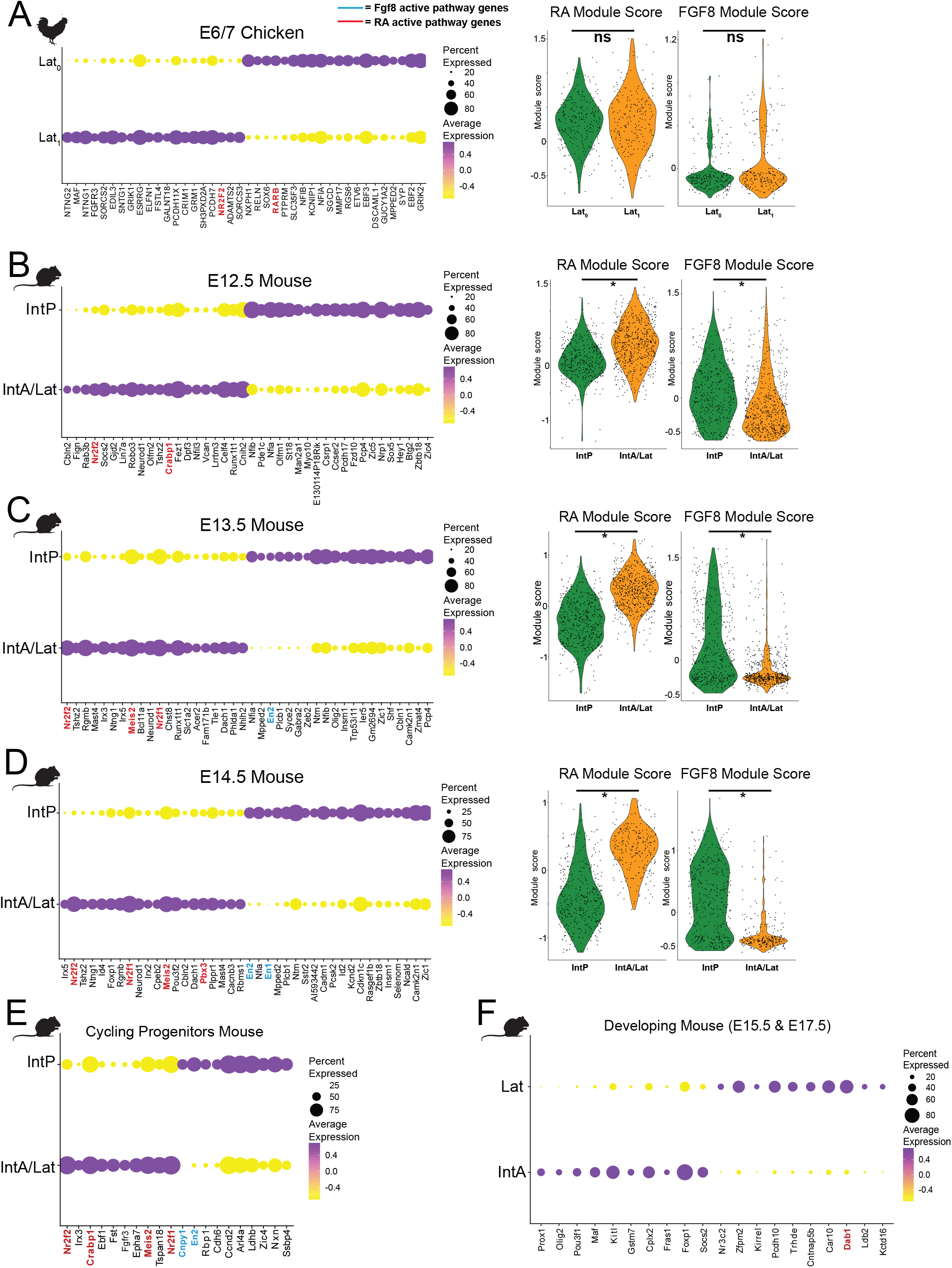
Chicken cerebellar nuclei lack mammalian RA- and FGF8-associated programs. **(A)** Dot plot of differentially expressed genes between Lat_0_ and Lat_1_ in E6/7 chicken (left), with retinoic acid (RA) and fibroblast growth factor 8 (FGF8) module scores across Lat_0_ and Lat_1_ (right). **(B-E)** Missing RA and FGF8 signatures in chicken are unlikely to be the result of a failure to match up chicken and mouse developmental ages correctly, as these pathways are active across a long time window in mice. (**B**) Dot plot of differentially expressed genes between IntP and IntA/Lat in E12.5 mouse (left), with RA and FGF8 module scores across IntP and IntA/Lat (right). **(C)** Similar to **(B)** for E13.5 mouse. **(D)** Similar to (**B)** for E14.5 mouse. **(E)** Dot plot of most differentially expressed genes between putative IntP and IntA/Lat cycling progenitors. **(F)** Dot plot of differentially expressed genes between IntA and Lat (lateral nucleus) in E15.5 and E17.5 mouse, with *Dab1* highlighted.

**Extended Data Figure 9.**
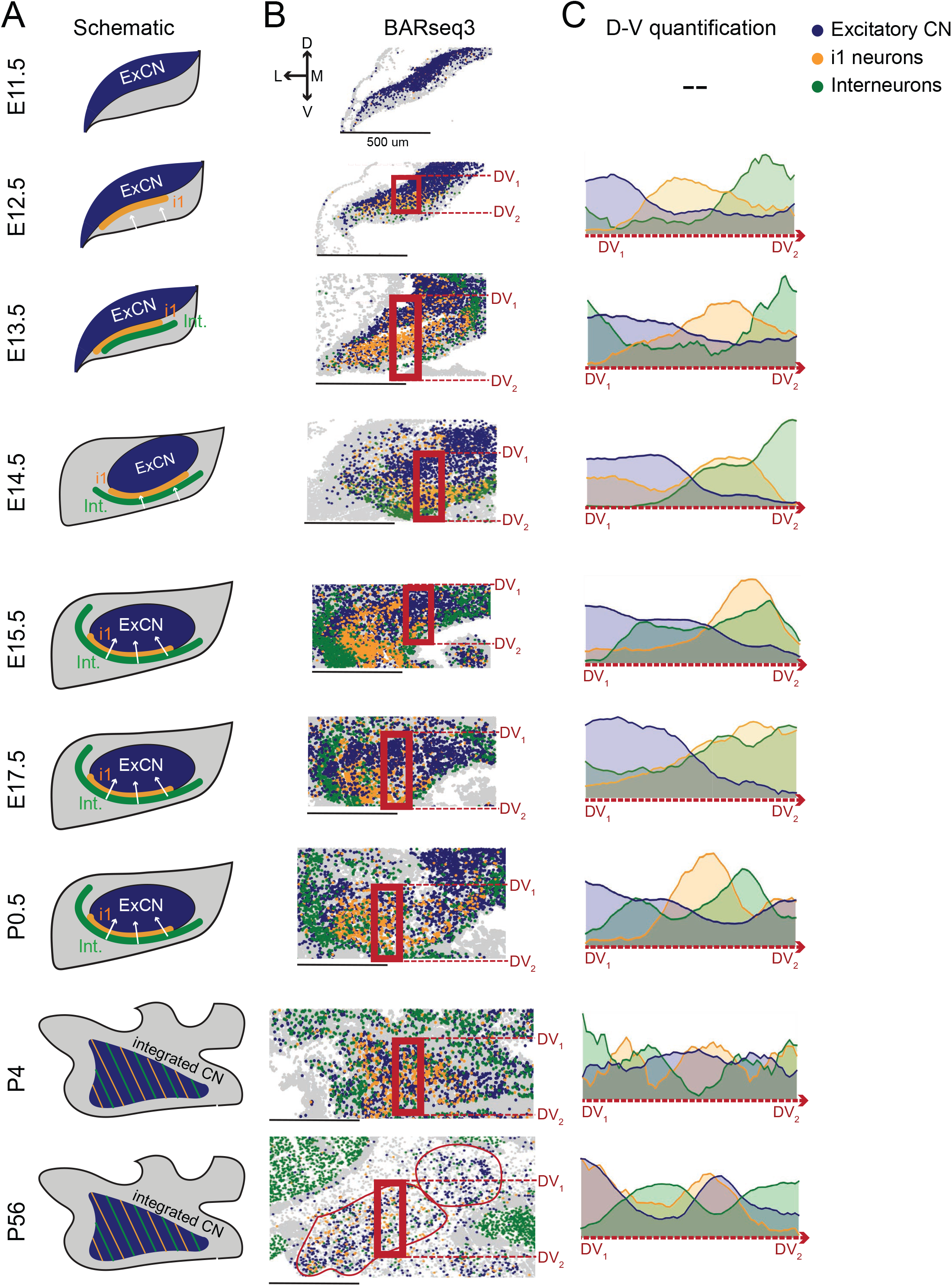
Inhibitory cerebellar nuclei neurons progressively integrate into excitatory territories. **(A)** Schematic of excitatory cerebellar nuclei neurons (blue), inhibitory i1 neurons (orange), and interneurons (green) across developmental timepoints. **(B)** BARseq3 data showing excitatory cerebellar nuclei neurons (blue), inhibitory i1 neurons (orange), and interneurons (green) across developmental timepoints. **(C)** Quantification of the distribution of cell type abundances from **(B)** across the dorsoventral axis shown in **(B)**.

**Extended Data Figure 10.**
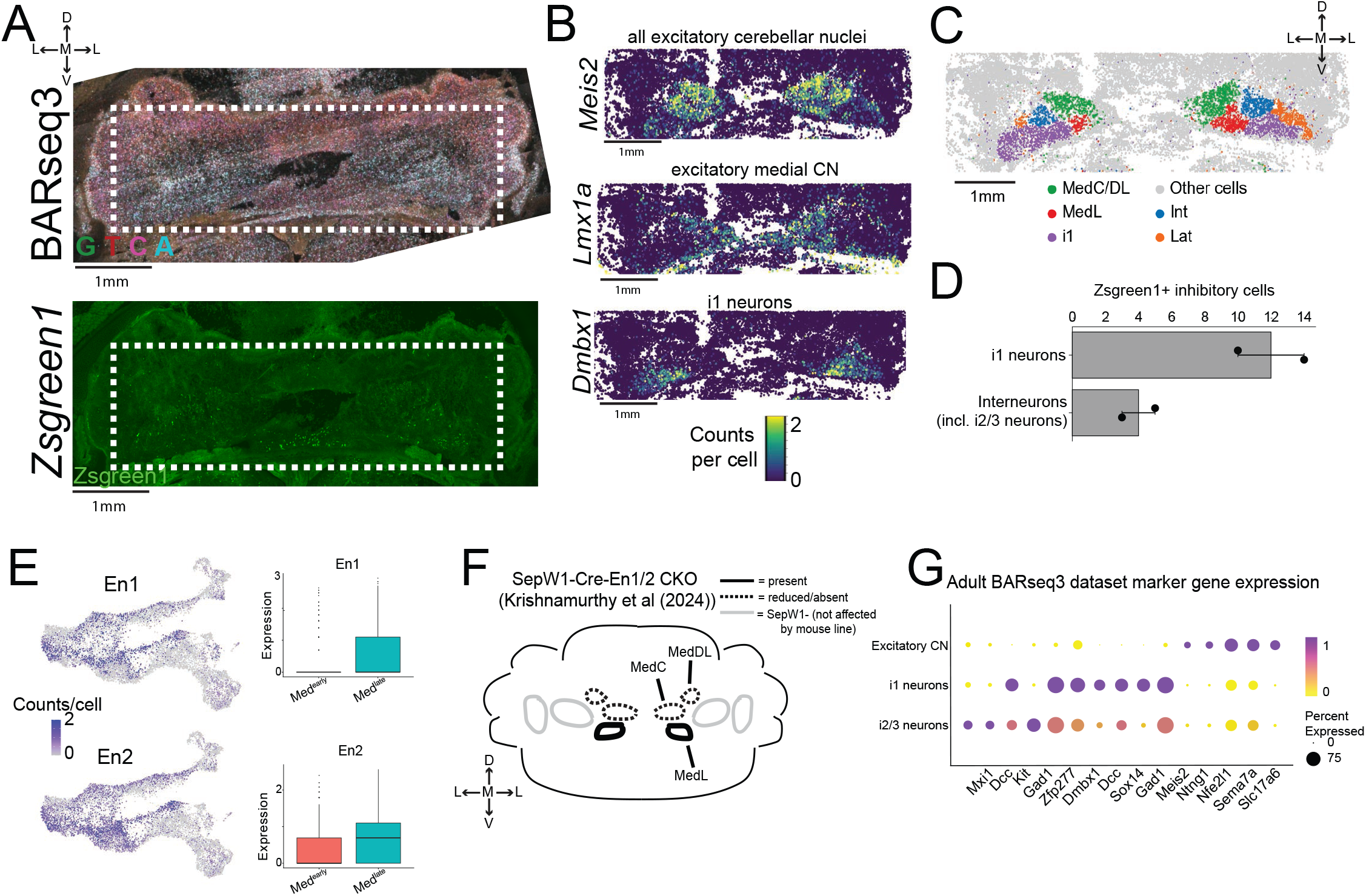
Med^early^ vs. Med^late^ population differences and BARseq3 of Robo3-CreERT2 x Ai6 mice. **(A)** Raw BARseq3 sequencing round 1 signal (top) and ZsGreen1 signal (bottom) in a Robo3-CreERT2 x Ai6 mouse at E18.5 after induction at E10.5, E11.5 and E12.5. White box indicates regions in which genes were decoded from BARseq3 analysis. **(B)** Expression of excitatory cerebellar nuclei marker *Meis2*, medial cerebellar nuclei marker *Lmx1a*, and i1 marker *Dmbx1* in ZsGreen1-labeled cells in the medial nucleus. **(C)** Spatial clustering of cerebellar nucleus neurons in Robo3-CreERT2 x Ai6 BARseq3 sections. **(D)** Number of i1 and i2/3 neurons labeled by ZsGreen1 in Robo3-CreERT2 x Ai6 BARseq3 sections. N=2 **(E)** Expression of *En1* and *En2* in the excitatory cerebellar nucleus UMAP embedding (left) and in violin plots (right), showing enrichment in Med^late^. **(F)** Summary of data showing in ^33^, where SepW1-Cre-En1/2 cKO resulted in loss of all medial cerebellar nuclei except for the MedL and part of the MedC. **(G)** Marker gene expression of populations shown in **Fig 6I,J**.

## SUPPLEMENTARY INFORMATION

### Supplementary Figures

**Supplementary Figure 1.**
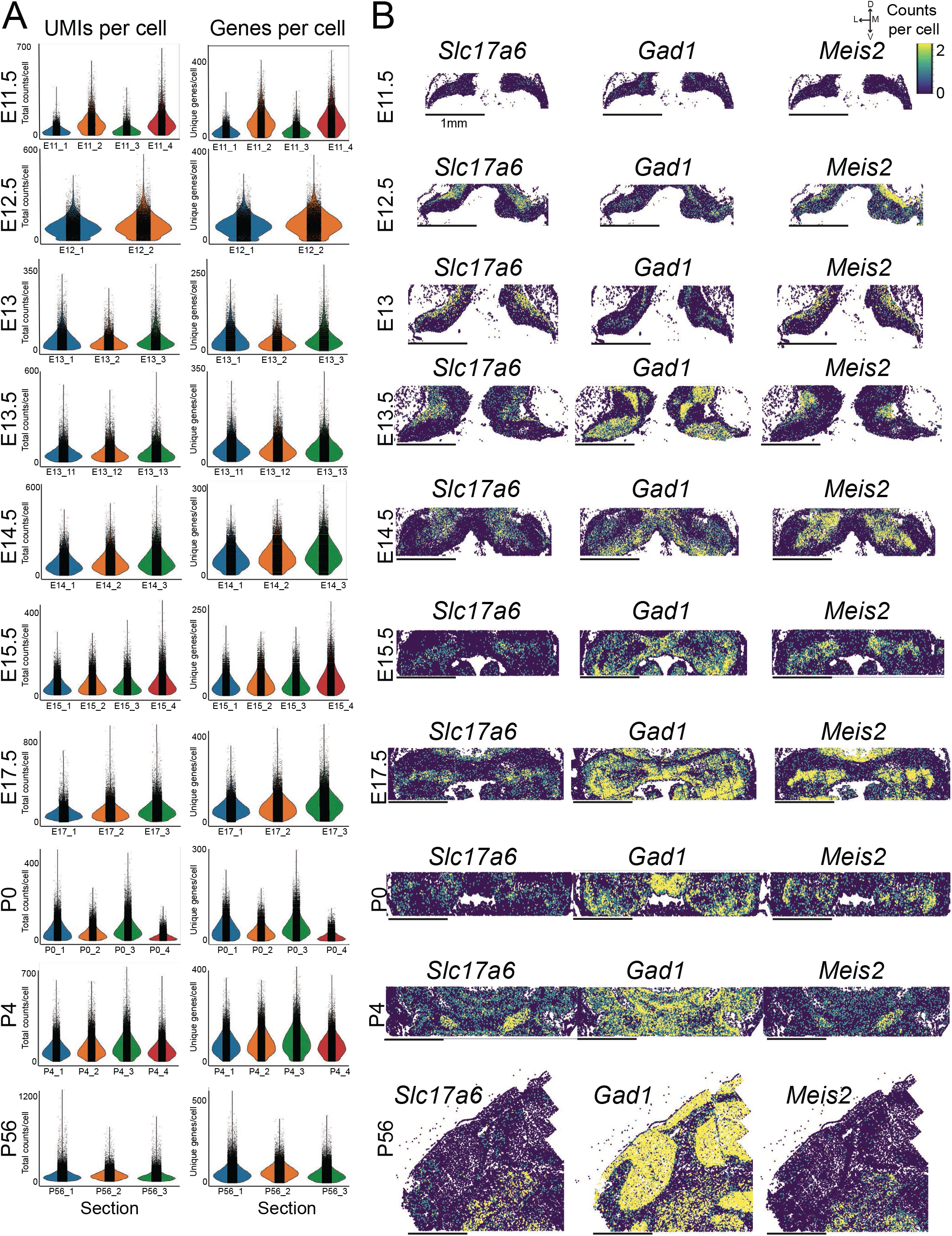
BARseq3 dataset quality control. **(A)** Number of UMIs per cell and number of unique genes per cell in each section for E11.5, E12.5, E13, E13.5, E14.5, E15.5, E17.5, P0, P4, and P56. **(B)** Spatial expression of *Slc17a6*, *Gad1*, and *Meis2* for one representative slice from each BARseq3 timepoint.

**Supplementary Figure 2.**
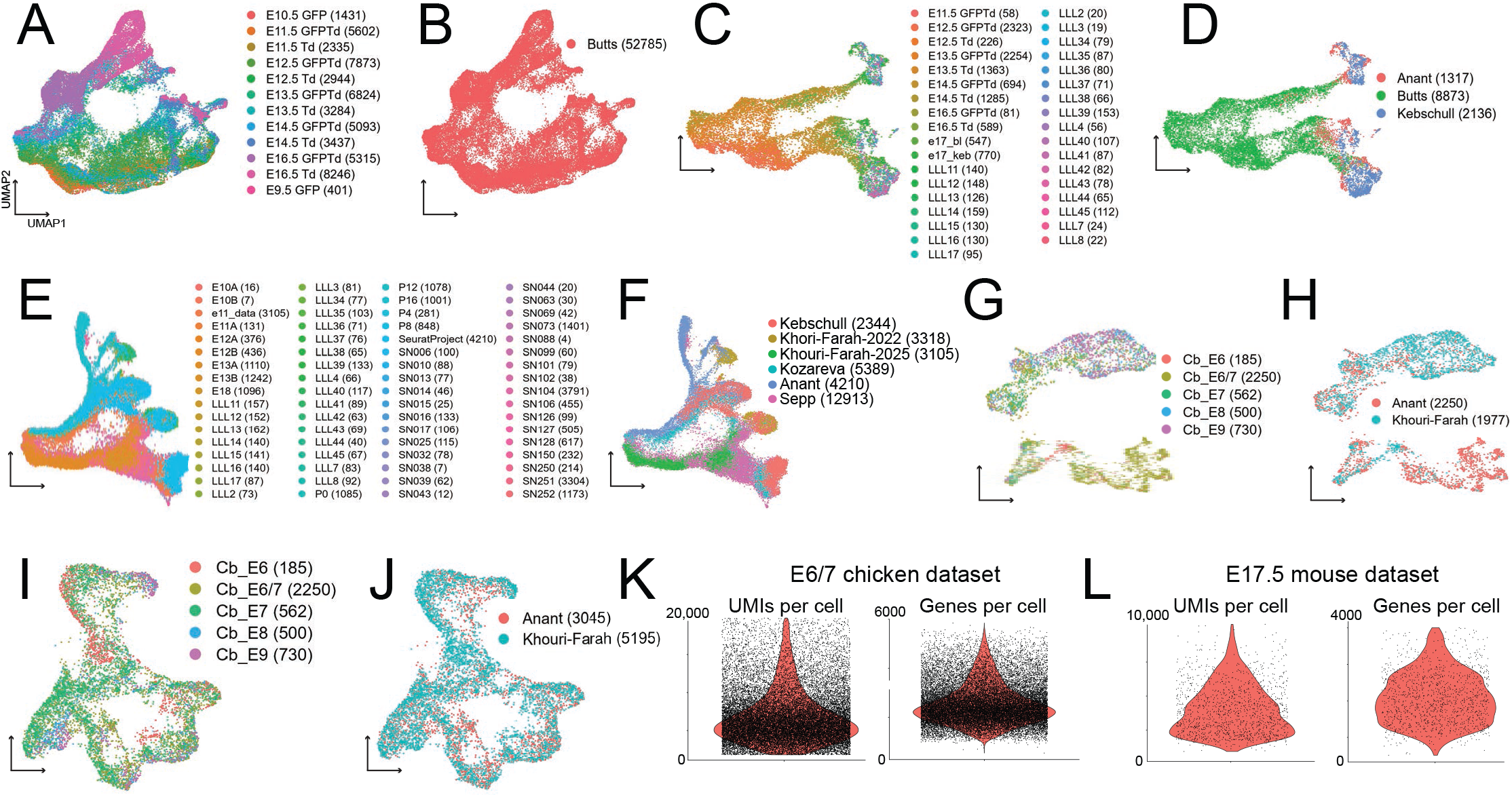
scRNAseq dataset integration and quality control. **(A,B)** UMAP embedding of the mouse upper rhombic lip Atoh1-lineage scRNAseq dataset, colored by **(A)** sample and **(B)** collector. Cell numbers contributed by each are given in parenthesis. (C,D) UMAP embedding of the integrated mouse excitatory cerebellar nuclei scRNAseq dataset, colored by **(C)** sample and **(D)** collector. **(E,F)** UMAP embedding of the integrated mouse cerebellar inhibitory neuron development scRNAseq dataset, colored by **(E)** sample and **(F)** collector. **(G,H)** UMAP embedding of the integrated chicken cerebellar excitatory neuron development scRNAseq dataset, colored by **(G)** sample and **(H)** collector. **(I,J)** UMAP embedding of the integrated chicken cerebellar inhibitory neuron development scRNAseq dataset, colored by **(I)** sample and **(J)** collector. **(K,L)** Number of UMIs per cell and number of unique genes per cell of the **(K)** E6/7 chicken and **(L)** E17.5 mouse datasets generated as part of this study.

**Supplementary Figure 3.**
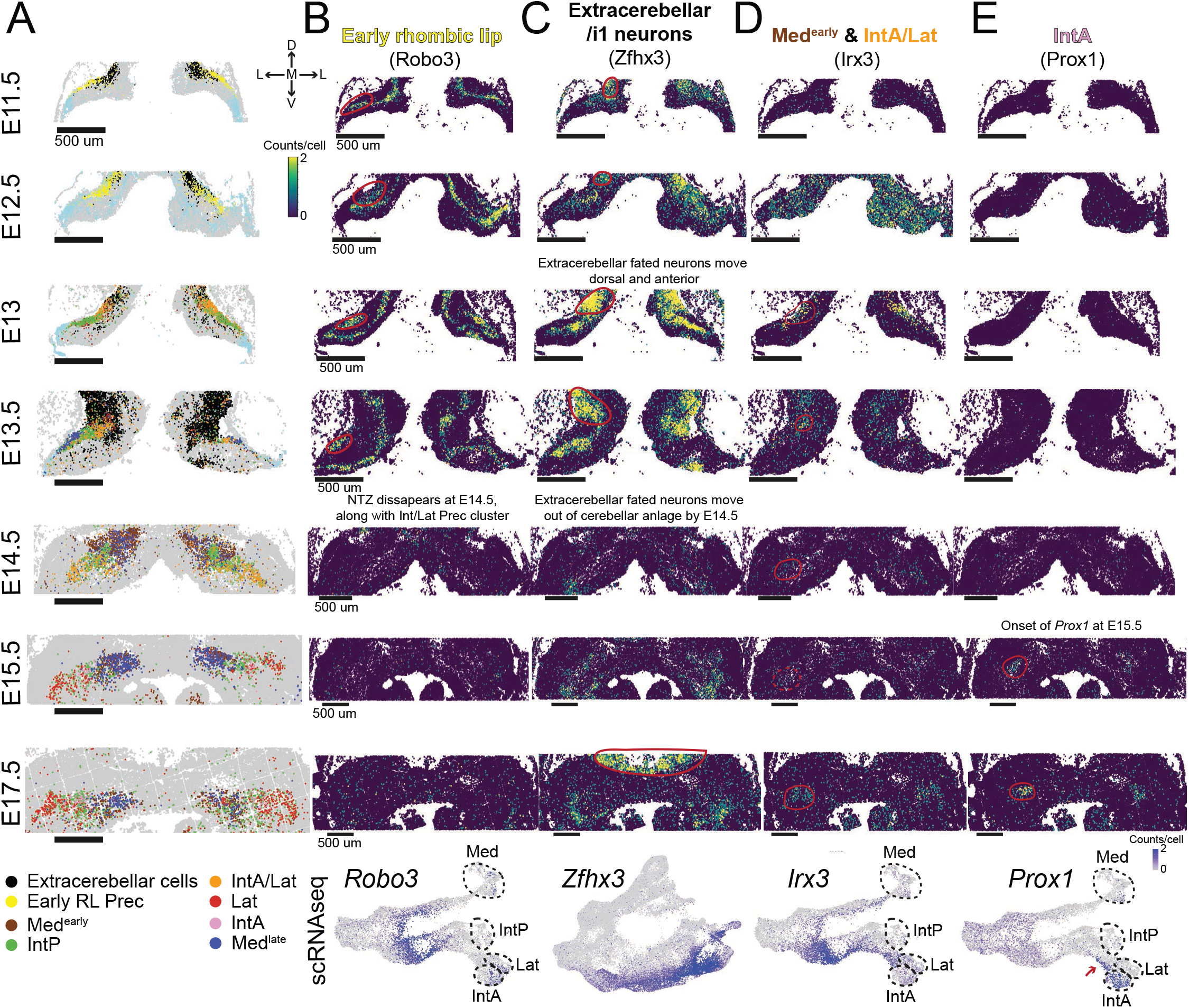
Spatial marker gene expression for early rhombic lip, extracerebellar neurons, Med^early^, IntA/Lat, and IntA cell types. BARseq3 datasets showing **(A)** excitatory cerebellar nuclei cell types across timepoints, **(B)** early rhombic lip *Robo3* expression (and inhibitory projection neurons), **(C)** extracerebellar/i1 neuron marker *Zfhx3*, **(D)** Med^early^ and IntA/Lat marker *Irx3*, and **(E)** IntA marker *Prox1*. The last row of each column contains scRNAseq expression of each gene. Red circles signify the location of the indicated cell type.

**Supplementary Figure 4.**
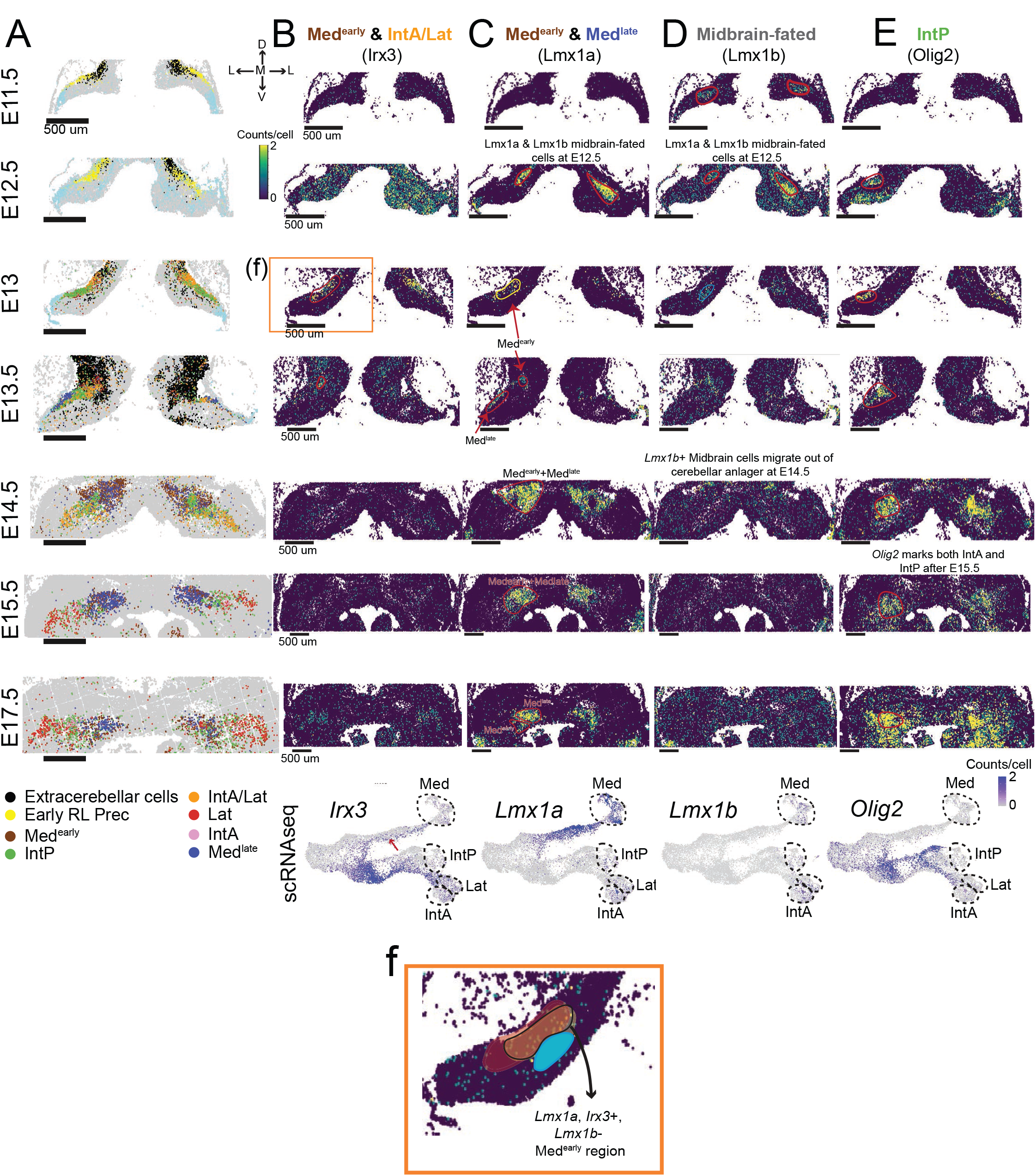
Spatial marker gene expression for Med^early^, IntA/Lat, Med^late^, midbrain-fated, and IntP cell types. BARseq3 datasets showing **(A)** excitatory cerebellar nuclei cell types across timepoints, **(B)** Med^early^ and IntA/Lat marker *Irx3*, **(C)** Med^early^ marker *Lmx1a*, **(D)** a *Lmx1a*+ population fated for the midbrain specifically, which is also marked by *Lmx1b* ^99^, and **(E)** IntP marker *Olig2*. The last row of each column contains scRNAseq expression of each gene. **(F)** Med^early^ is *Lmx1a*-positive (yellow) and *Irx3*-positive (red), but *Lmx1b*-negative (blue), shown in the region demarcated in black, and therefore does not correspond to the canonical medial nucleus or any extracerebellar-fated populations. Red circles signify the location of the indicated cell type.

**Supplementary Figure 5.**
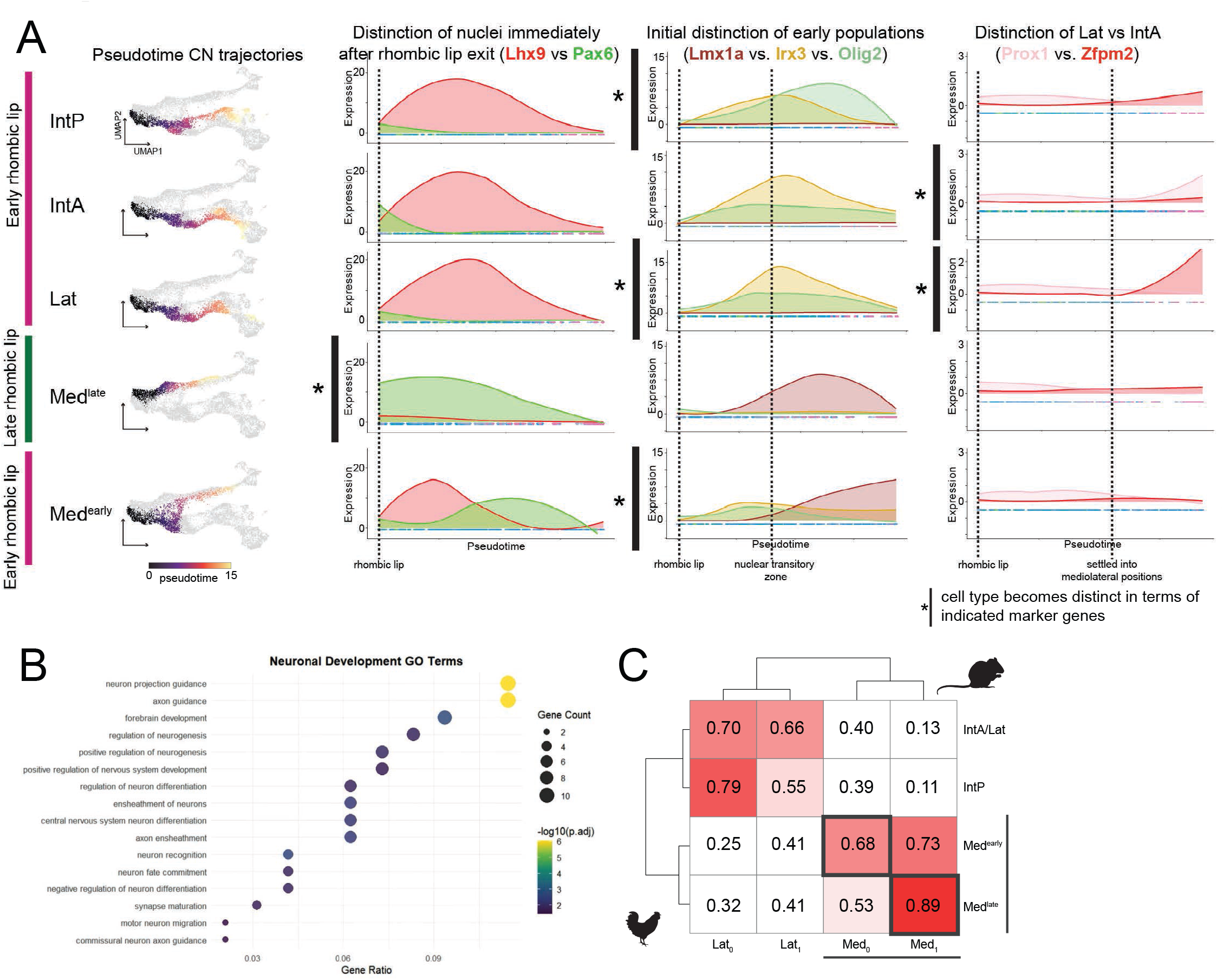
Pseudotime plot for each excitatory cerebellar nucleus trajectory across development. **(A)** Pseudotime plot based on scRNAseq depicting expression of *Lhx9* and *Pax6* at the rhombic lip, which distinguish Med^late^ from the other nuclei; *Lmx1a*, *Irx3*, and *Olig2*, which distinguish Med^early^, IntA/Lat, and IntP at the nuclear transitory zone; and Lat and IntA, which are distinguished by *Zfpm2* and *Prox1,* respectively, after nuclei settle into their mature mediolateral arrangement. (**B**) Gene ontology analysis of differentially expressed genes between Med^early^ and Med^late^, showing enrichment for axon guidance genes. (**C**) Correlation of mouse Med^early^ and Med^late^ populations with distinct medial populations in chicken during development.

**Supplementary Figure 6.**
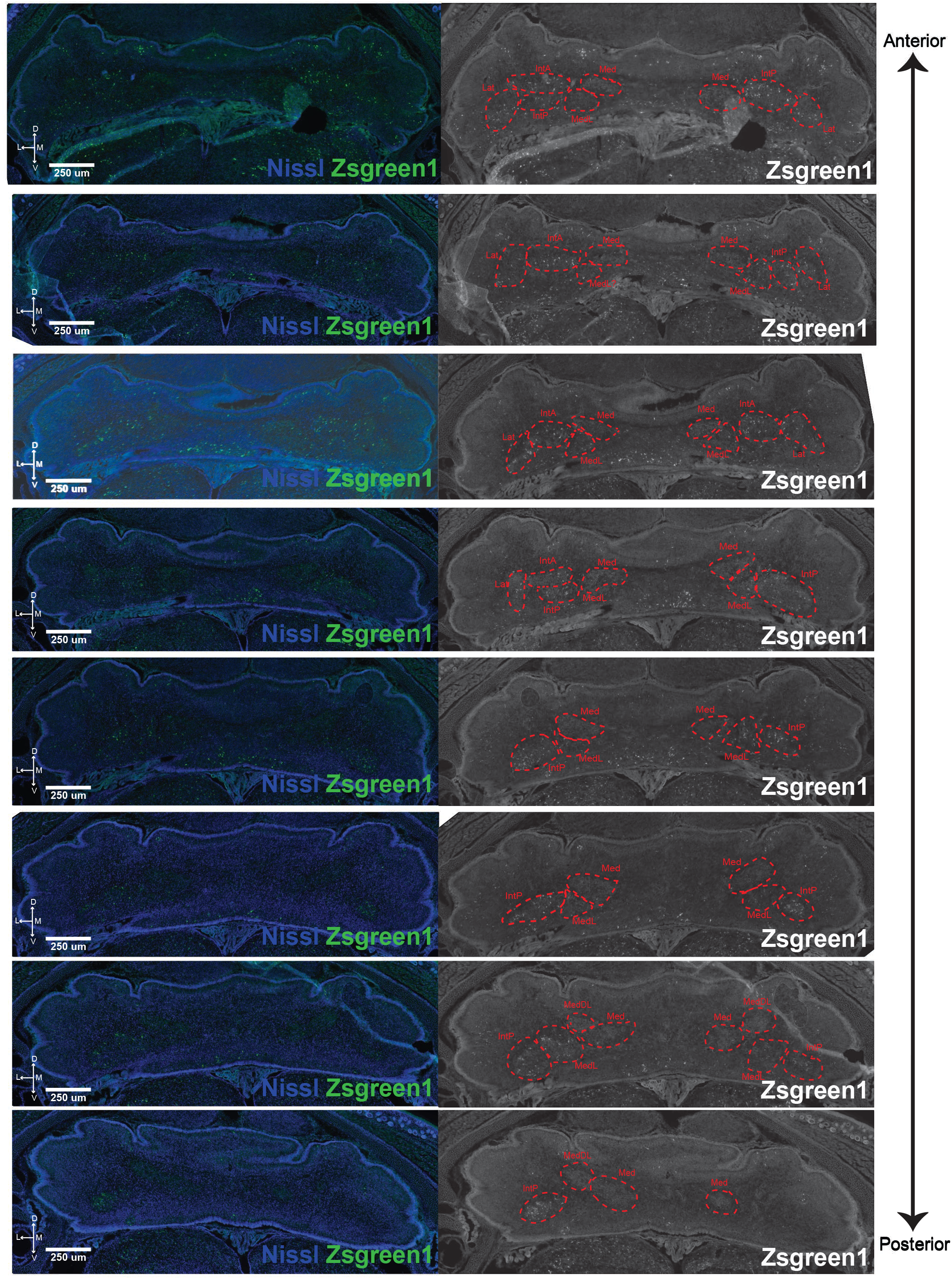
Robo3-CreERT2 x Ai6 mice label lateral, interposed, and MedL cerebellar (sub)nuclei. Representative sections of a Robo3-CreERT2 x Ai6 mouse at E18.5 after tamoxifen induction at E10.5, E11.5 and E12.5, showing Nissl staining and ZsGreen1 (left) and ZsGreen1 alone (right) along the anterior-posterior axis of the cerebellar nuclei. N=2 embryos.

## Notes

### Competing Interest Statement

The authors have declared no competing interest.

### Summary of Updates

In this revised manuscript, we updated the abstract and introduction.

## REFERENCES

1. Chakraborty, M. & Jarvis, E. D. Brain evolution by brain pathway duplication. Philos. Trans. R. Soc. Lond. B Biol. Sci. 370, (2015).

2. Tosches, M. A. Developmental and genetic mechanisms of neural circuit evolution. Dev. Biol. 431, 16–25 (2017).

3. Kebschull, J. M. et al. Cerebellar nuclei evolved by repeatedly duplicating a conserved cell-type set. Science 370, (2020).

4. Grillner, S. & Robertson, B. The basal ganglia over 500 million years. Curr. Biol. 26, R1088–R1100 (2016).

5. Frangeul, L. et al. A cross-modal genetic framework for the development and plasticity of sensory pathways. Nature 538, 96–98 (2016).

6. Arendt, D. The evolution of cell types in animals: emerging principles from molecular studies. Nat. Rev. Genet. 9, 868–882 (2008).

7. Kebschull, J. M. et al. Cerebellum lecture: The cerebellar nuclei-core of the cerebellum. Cerebellum 23, 620–677 (2024).

8. Anant, M. M.-G., Faltine-Gonzalez, D. Z., Lacava, S. A. & Kebschull, J. M. Principles of brain region evolution: Insights from the cerebellar nuclei. Annu. Rev. Neurosci. 49, 39–59 (2026).

9. The Cerebellum of Non-Mammalian Vertebrates. in Evolution of Nervous Systems, 2nd Edition, Vol I. 373–385 (JElsevier, 2017).

10. Chambers, W. W. & Sprague, J. M. Functional localization in the cerebellum. II. Somatotopic organization in cortex and nuclei. AMA Arch. Neurol. Psychiatry 74, 653–680 (1955).

11. Gao, Z. et al. A cortico-cerebellar loop for motor planning. Nature 563, 113–116 (2018).

12. De Schutter, E. Fallacies of mice experiments. Neuroinformatics 17, 181–183 (2019).

13. Fujita, H., Kodama, T. & du Lac, S. Modular output circuits of the fastigial nucleus for diverse motor and nonmotor functions of the cerebellar vermis. Elife 9, e58613 (2020).

14. Helgers, S. O. A. et al. Lesion of the fastigial nucleus in juvenile rats deteriorates rat behavior in adulthood, accompanied by altered neuronal activity in the medial prefrontal cortex. Neuroscience 442, 29–40 (2020).

15. Streng, M. L., Tetzlaff, M. R. & Krook-Magnuson, E. Distinct fastigial output channels and their impact on temporal lobe seizures. J. Neurosci. 41, 10091–10107 (2021).

16. Urrutia Desmaison, J. D., et al. Cerebellar control of fear learning via the cerebellar nuclei-Multiple pathways, multiple mechanisms? Front. Syst. Neurosci. 17, 1176668 (2023).

17. Qi, H., et al. BARseq3: a modular system for integrating spatial multi-omics and cellular barcoding in single cells. bioRxivorg (2026) doi:10.64898/2026.05.13.724900.

18. Machold, R. & Fishell, G. Math1 is expressed in temporally discrete pools of cerebellar rhombic-lip neural progenitors. Neuron 48, 17–24 (2005).

19. Sekerková, G., Ilijic, E. & Mugnaini, E. Time of origin of unipolar brush cells in the rat cerebellum as observed by prenatal bromodeoxyuridine labeling. Neuroscience 127, 845–858 (2004).

20. Wang, V. Y., Rose, M. F. & Zoghbi, H. Y. Math1 expression redefines the rhombic lip derivatives and reveals novel lineages within the brainstem and cerebellum. Neuron 48, 31–43 (2005).

21. Leto, K. et al. Consensus paper: Cerebellar development. Cerebellum 15, 789–828 (2016).

22. Leto, K. & Rossi, F. Specification and differentiation of cerebellar GABAergic neurons. Cerebellum 11, 434–435 (2012).

23. Sudarov, A. et al. Ascl1 genetics reveals insights into cerebellum local circuit assembly. J. Neurosci. 31, 11055–11069 (2011).

24. Engelkamp, D., Rashbass, P., Seawright, A. & van Heyningen, V. Role of Pax6 in development of the cerebellar system. Development 126, 3585–3596 (1999).

25. Bulfone, A. et al. Expression pattern of the Tbr2 (Eomesodermin) gene during mouse and chick brain development. Mech. Dev. 84, 133–138 (1999).

26. Fink, A. J. et al. Development of the deep cerebellar nuclei: transcription factors and cell migration from the rhombic lip. J. Neurosci. 26, 3066–3076 (2006).

27. Khouri-Farah, N., Guo, Q., Perry, T. A., Dussault, R. & Li, J. Y. H. FOXP genes regulate Purkinje cell diversity and cerebellar morphogenesis. Nat. Neurosci. 28, 2022–2033 (2025).

28. Khouri-Farah, N., Guo, Q., Morgan, K., Shin, J. & Li, J. Y. H. Integrated single-cell transcriptomic and epigenetic study of cell state transition and lineage commitment in embryonic mouse cerebellum. Sci. Adv. 8, eabl9156 (2022).

29. Kozareva, V. et al. A transcriptomic atlas of mouse cerebellar cortex comprehensively defines cell types. Nature 598, 214–219 (2021).

30. Sepp, M. et al. Cellular development and evolution of the mammalian cerebellum. Nature 625, 788–796 (2024).

31. Schneider, B. F. & Norton, S. Equivalent ages in rat, mouse and chick embryos. Teratology 19, 273–278 (1979).

32. Wizeman, J. W., Guo, Q., Wilion, E. M. & Li, J. Y. Specification of diverse cell types during early neurogenesis of the mouse cerebellum. Elife 8, e42388 (2019).

33. Krishnamurthy, A. et al. Engrailed transcription factors direct excitatory cerebellar neuron diversity and survival. Development 151, dev202502 (2024).

34. Akçay, V., et al. Conserved cerebellar rhombic lip compartmentalization and *Eomes* regulatory networks govern unipolar brush cell development. bioRxiv (2026) doi:10.64898/2026.07.27.740783.

35. Schilling, K. Revisiting the development of cerebellar inhibitory interneurons in the light of single-cell genetic analyses. Histochem. Cell Biol. 161, 5–27 (2024).

36. Green, M. J. & Wingate, R. J. T. Developmental origins of diversity in cerebellar output nuclei. Neural Dev. 9, 1 (2014).

37. Wingate, R. J. The rhombic lip and early cerebellar development. Curr. Opin. Neurobiol. 11, 82–88 (2001).

38. Wilson, L. J. & Wingate, R. J. T. Temporal identity transition in the avian cerebellar rhombic lip. Dev. Biol. 297, 508–521 (2006).

39. Butts, T., Green, M. J. & Wingate, R. J. T. Development of the cerebellum: simple steps to make a “little brain.” Development 141, 4031–4041 (2014).

40. Butts, J. C. et al. A single-cell transcriptomic map of the developing Atoh1 lineage identifies neural fate decisions and neuronal diversity in the hindbrain. Dev. Cell 59, 2171–2188.e7 (2024).

41. Martí-Clua, J. Times of neuron origin and neurogenetic gradients in mice Purkinje cells and deep cerebellar nuclei neurons during the development of the cerebellum. A review. Tissue Cell 78, 101897 (2022).

42. Casoni, F. et al. A spatial-temporal map of glutamatergic neurogenesis in the murine embryonic cerebellar nuclei uncovers a high degree of cellular heterogeneity. J. Anat. 245, 560–571 (2024).

43. Clark, B. S. et al. Single-cell RNA-seq analysis of retinal development identifies NFI factors as regulating mitotic exit and late-born cell specification. Neuron 102, 1111–1126.e5 (2019).

44. Tang, K. et al. COUP-TFs regulate eye development by controlling factors essential for optic vesicle morphogenesis. Development 137, 725–734 (2010).

45. Lyu, P. et al. Gene regulatory networks controlling temporal patterning, neurogenesis, and cell-fate specification in mammalian retina. Cell Rep. 37, 109994 (2021).

46. Sagner, A. et al. A shared transcriptional code orchestrates temporal patterning of the central nervous system. PLoS Biol. 19, e3001450 (2021).

47. da Silva, S. & Cepko, C. L. Fgf8 expression and degradation of retinoic acid are required for patterning a high-acuity area in the retina. Dev. Cell 42, 68–81.e6 (2017).

48. Cunningham, T. J. et al. Retinoic acid activity in undifferentiated neural progenitors is sufficient to fulfill its role in restricting Fgf8 expression for somitogenesis. PLoS One 10, e0137894 (2015).

49. Lewandoski, M. & Mackem, S. Limb development: the rise and fall of retinoic acid. Curr. Biol. 19, R558–61 (2009).

50. Diez del Corral, R., et al. Opposing FGF and retinoid pathways control ventral neural pattern, neuronal differentiation, and segmentation during body axis extension. Neuron 40, 65–79 (2003).

51. Zhang, J., Smith, D., Yamamoto, M., Ma, L. & McCaffery, P. The meninges is a source of retinoic acid for the late-developing hindbrain. J. Neurosci. 23, 7610–7620 (2003).

52. Chi, C. L., Martinez, S., Wurst, W. & Martin, G. R. The isthmic organizer signal FGF8 is required for cell survival in the prospective midbrain and cerebellum. Development 130, 2633–2644 (2003).

53. Harada, H., Sato, T. & Nakamura, H. Fgf8 signaling for development of the midbrain and hindbrain. Dev. Growth Differ. 58, 437–445 (2016).

54. Wilson, L. J., Myat, A., Sharma, A., Maden, M. & Wingate, R. J. T. Retinoic acid is a potential dorsalising signal in the late embryonic chick hindbrain. BMC Dev. Biol. 7, 138 (2007).

55. Sinagra, M. et al. Glutamatergic cerebellar granule neurons synthesize and secrete reelin in vitro. Neuron Glia Biol. 4, 189–196 (2008).

56. Utsunomiya-Tate, N. et al. Reelin molecules assemble together to form a large protein complex, which is inhibited by the function-blocking CR-50 antibody. Proc. Natl. Acad. Sci. U. S. A. 97, 9729–9734 (2000).

57. Buckley, P. F. Lower number of cerebellar Purkinje neurons in psychosis is associated with reduced reelin expression. Year B. Psychiatry Appl. Ment. Health 2011, 366–368 (2011).

58. Gao, Z. & Godbout, R. Reelin-Disabled-1 signaling in neuronal migration: splicing takes the stage. Cell. Mol. Life Sci. 70, 2319–2329 (2013).

59. Howell, B. W., Herrick, T. M. & Cooper, J. A. Reelin-induced tyrosine [corrected] phosphorylation of disabled 1 during neuronal positioning. Genes Dev. 13, 643–648 (1999).

60. Caviness, V. S., Jr & Sidman, R. L. Olfactory structures of the forebrain in the reeler mutant mouse. J. Comp. Neurol. 145, 85–104 (1972).

61. Goffinet, A. M., So, K. F., Yamamoto, M., Edwards, M. & Caviness, V. S., Jr. Architectonic and hodological organization of the cerebellum in reeler mutant mice. Brain Res. 318, 263–276 (1984).

62. Goffinet, A. M. The embryonic development of the cerebellum in normal and reeler mutant mice. Anat. Embryol. (Berl.) 168, 73–86 (1983).

63. Li, H. et al. Classifying Drosophila olfactory projection neuron subtypes by single-cell RNA sequencing. Cell 171, 1206–1220.e22 (2017).

64. Leto, K., Carletti, B., Williams, I. M., Magrassi, L. & Rossi, F. Different types of cerebellar GABAergic interneurons originate from a common pool of multipotent progenitor cells. J. Neurosci. 26, 11682–11694 (2006).

65. Prekop, H.-T. et al. Sox14 is required for a specific subset of cerebello-olivary projections. J. Neurosci. 38, 9539–9550 (2018).

66. Seto, Y. et al. Temporal identity transition from Purkinje cell progenitors to GABAergic interneuron progenitors in the cerebellum. Nat. Commun. 5, 3337 (2014).

67. Bagnall, M. W. et al. Glycinergic projection neurons of the cerebellum. J. Neurosci. 29, 10104–10110 (2009).

68. Lee, A. S. et al. Cerebellar output neurons can impair non-motor behaviors by altering development of extracerebellar connectivity. Nat. Commun. 16, 1858 (2025).

69. Bengtsson, F., Rasmussen, A. & Hesslow, G. Feedback control in the olivocerebellar loop. in Handbook of the Cerebellum and Cerebellar Disorders 1215–1238 (Springer International Publishing, Cham, 2022).

70. Garwicz, M. Olivocerebellar pathway. in Encyclopedia of Computational Neuroscience 2532–2536 (Springer New York, New York, NY, 2022).

71. Lang, E. J. et al. The roles of the olivocerebellar pathway in motor learning and motor control. A consensus paper. Cerebellum 16, 230–252 (2017).

72. Wang, X. et al. Excitatory nucleo-olivary pathway shapes cerebellar outputs for motor control. Nat. Neurosci. 26, 1394–1406 (2023).

73. Agirman, G., Broix, L. & Nguyen, L. Cerebral cortex development: an outside-in perspective. FEBS Lett. 591, 3978–3992 (2017).

74. Isko, E. C. et al. Specific expansion of motor cortical projections in a singing mouse. Nature 655, 438–446 (2026).

75. Tyssowski, K. M. et al. Evolutionary expansion of the corticospinal system is linked to dexterity in Peromyscus mice. bioRxivorg (2025) doi:10.1101/2025.10.16.682851.

76. Kaas, J. H. The evolution of neocortex in primates. Prog. Brain Res. 195, 91–102 (2012).

77. Zaremba, B. et al. Developmental origins and evolution of pallial cell types and structures in birds. Science 387, eadp5182 (2025).

78. Hain, D. et al. Molecular diversity and evolution of neuron types in the amniote brain. Science 377, eabp8202 (2022).

79. Tosches, M. A. et al. Evolution of pallium, hippocampus, and cortical cell types revealed by single-cell transcriptomics in reptiles. Science 360, 881–888 (2018).

80. Colquitt, B. M., Merullo, D. P., Konopka, G., Roberts, T. F. & Brainard, M. S. Cellular transcriptomics reveals evolutionary identities of songbird vocal circuits. Science 371, (2021).

81. Woych, J. et al. Cell-type profiling in salamanders identifies innovations in vertebrate forebrain evolution. Science 377, eabp9186 (2022).

82. Voogd, J., Shinoda, Y., Ruigrok, T. J. H. & Sugihara, I. Cerebellar nuclei and the inferior olivary nuclei: Organization and connections. in Handbook of the Cerebellum and Cerebellar Disorders 497–557 (Springer International Publishing, Cham, 2022).

83. Willett, R. T. et al. Cerebellar nuclei excitatory neurons regulate developmental scaling of presynaptic Purkinje cell number and organ growth. Elife 8, (2019).

84. Hamburger, V. & Hamilton, H. L. A series of normal stages in the development of the chick embryo. Dev. Dyn. 195, 231–272 (1992).

85. Chou, H.-H. Shared probe design and existing microarray reanalysis using PICKY. BMC Bioinformatics 11, 196 (2010).

86. Schmidt, T. L. et al. Scalable amplification of strand subsets from chip-synthesized oligonucleotide libraries. Nat. Commun. 6, 8634 (2015).

87. Bryan, W. P. & Byrne, R. H. A calcium chloride solution, dry-ice, low temperature bath. J. Chem. Educ. 47, 361 (1970).

88. Tangeman, J. A., Charris Dominguez, C. M., Bendezu-Sayas, S. & Del Rio-Tsonis, K. Nuclei isolation from ocular tissues of the embryonic chicken for single-nucleus profiling. Methods Mol. Biol. 2848, 105–116 (2025).

89. Haghverdi, L., Lun, A. T. L., Morgan, M. D. & Marioni, J. C. Batch effects in single-cell RNA-sequencing data are corrected by matching mutual nearest neighbors. Nat. Biotechnol. 36, 421–427 (2018).

90. Hao, Y. et al. Integrated analysis of multimodal single-cell data. Cell 184, 3573–3587.e29 (2021).

91. Hafemeister, C. & Satija, R. Normalization and variance stabilization of single-cell RNA-seq data using regularized negative binomial regression. Genome Biol. 20, 296 (2019).

92. Friedman, J., Hastie, T. & Tibshirani, R. Regularization paths for generalized linear models via coordinate descent. J. Stat. Softw. 33, 1–22 (2010).

93. Wolf, F. A., Angerer, P. & Theis, F. J. SCANPY: large-scale single-cell gene expression data analysis. Genome Biol. 19, 15 (2018).

94. Korsunsky, I. et al. Fast, sensitive and accurate integration of single-cell data with Harmony. Nat. Methods 16, 1289–1296 (2019).

95. Palla, G. et al. Squidpy: a scalable framework for spatial omics analysis. Nat. Methods 19, 171–178 (2022).

96. Crow, M., Paul, A., Ballouz, S., Huang, Z. J. & Gillis, J. Characterizing the replicability of cell types defined by single cell RNA-sequencing data using MetaNeighbor. Nat. Commun. 9, 884 (2018).

97. Paxinos, G. & Franklin, K. B. J. Paxinos and Franklin’s the Mouse Brain in Stereotaxic Coordinates. (Academic Press, San Diego, CA, 2019).

98. Sicotte, N. L. et al. Diffusion tensor MRI shows abnormal brainstem crossing fibers associated with ROBO3 mutations. Neurology 67, 519–521 (2006).

99. Millen, K. J., Steshina, E. Y., Iskusnykh, I. Y. & Chizhikov, V. V. Transformation of the cerebellum into more ventral brainstem fates causes cerebellar agenesis in the absence of Ptf1a function. Proc. Natl. Acad. Sci. U. S. A. 111, E1777–86 (2014).

